# Autonomous emergence of connectivity assemblies via spike triplet interactions

**DOI:** 10.1101/716001

**Authors:** Lisandro Montangie, Julijana Gjorgjieva

## Abstract

Non-random connectivity can emerge without structured external input driven by activity-dependent mechanisms of synaptic plasticity based on precise spiking patterns. Here we analyze the emergence of global structures in recurrent networks based on a triplet model of spike timing dependent plasticity (STDP) which depends on the interactions of three precisely-timed spikes and can describe plasticity experiments with varying spike frequency better than the classical pair-based STDP rule. We describe synaptic changes arising from emergent higher-order correlations, and investigate their influence on different connectivity motifs in the network. Our motif expansion framework reveals novel motif structures under the triplet STDP rule, which support the formation of bidirectional connections and loops in contrast to the classical pair-based STDP rule. Therefore, triplet STDP drives the spontaneous emergence of self-connected groups of neurons, or assemblies, proposed to represent functional units in neural circuits. Assembly formation has often been associated with plasticity driven by firing rates or external stimuli. We propose that assembly structure can emerge without the need for externally patterned inputs or assuming a symmetric pair-based STDP rule commonly assumed in previous studies. The emergence of non-random network structure under triplet STDP occurs through internally-generated higher-order correlations, which are ubiquitous in natural stimuli and neuronal spiking activity, and important for coding. We further demonstrate how neuromodulatory mechanisms that modulate the shape of triplet STDP or the synaptic transmission function differentially promote connectivity motifs underlying the emergence of assemblies, and quantify the differences using graph theoretic measures.

## Introduction

The synaptic wiring between neurons – originally proposed as a mechanism for learning and memory – is sculpted by experience and has become a most relevant link between circuit structure and function [1]. The original formulation of Hebbian plasticity, whereby “cells that fire together, wire together” [2,3], fostered the concept of ‘cell assemblies’ [4], defined as groups of neurons that are repeatedly co-activated leading to the strengthening of synaptic connectivity between individual neurons. This has suggested that activity-dependent synaptic plasticity, including both long-term potentiation and long-term depression, is a key mechanism for the emergence of assemblies in the organization of neural circuits [5–7]. These interconnected groups of neurons have become an important target for many theories of neural computation and associative memory [8–11]. Recent technological developments that enable multiple neurons to be simultaneously recorded have provided the much needed physiological evidence of assembly organization [12–15]. For instance, synaptically connected neurons tend to receive more common input than would be expected by chance, [12,16–18] and cortical pyramidal neurons tend to be more strongly connected to neurons that share stimulus preference [13,19,20], providing evidence for clustered architecture. It has been proposed that this organization enables the cortex to intrinsically generate reverberating patterns of neural activity when representing different stimulus features [1,21]. Thus, neuronal assemblies can be interpreted as the building blocks of cortical microcircuits which are differentially recruited during distinct functions, such as the binding of different features of a sensory stimulus [7,17,22]. In addition to cortical circuits, neuronal assemblies have also been observed in the optic tectum (a structure homologous to the superior colliculus in mammals [23]) in the developing zebrafish larva [24–27]. Experiments in sensory deprived larvae have demonstrated that the basic structure of spontaneous activity and functional connectivity emerges without intact retinal inputs, suggesting that neuronal assemblies are intrinsically generated in the tectum and not just the product of correlated external inputs [25–27]. This raises the important question of what drives the emergence of these clustered structures, and whether patterned external input is necessary.

To understand the emergence of such non-random connectivity, a growing body of theoretical and computational work has been developed to link connectivity architecture to the coordinated spiking activity of neurons, especially in recurrent networks [28–41]. These studies can be divided into two classes: those that examine the influence of externally structured input on activity-dependent refinement [42–45], and those that investigate the autonomous emergence of non-random connectivity in the absence of patterned external input, purely driven by emergent network interactions [5,6,46]. Specifically, assemblies in recurrent networks can be imprinted based on internally-generated network interactions [6] or through rate-based plasticity where inputs with higher firing rates to subsets of neurons strengthen recurrent connections [47,48]; assemblies can also be initially determined by externally patterned input but maintained by internal correlations [49].

Despite this success, all of these studies have assumed pair-based models of STDP, which induce plasticity based on the precise timing and order of a pair of pre- and postsynaptic spikes [50,51]. Here, we consider a spike-based learning rule, “the triplet STDP model” [52], which uses sets of three spikes (triplets) to induce potentiation. More precisely, potentiation depends on the interval between the pre- and postsynaptic spikes, and on the timing of the previous postsynaptic spike. This triplet learning rule has been shown to explain a variety of synaptic plasticity data [53,54] significantly better than pair-based STDP [52]. We have previously shown a tight correspondence between the triplet STDP rule and the well-known Bienenstock-Cooper-Munro (BCM) synaptic learning rule, which maximizes the selectivity of the postsynaptic neuron, and thereby offers a possible explanation for experience-dependent cortical plasticity such as orientation and direction selectivity. In addition, triplet STDP can also induce selectivity for input patterns consisting of higher-order correlations (HOCs). HOCs have been experimentally measured in several brain areas [55], and shown to account for a substantial amount of information transfer in sensory cortex [55–58]. HOCs are also important for characterizing the firing of a postsynaptic neuron [59,60], circuit function and coding [61,62], and the synchronous firing and the distribution of activity in a neuronal pool [63–66]. Here we investigated the functional significance of such HOCs for shaping recurrent network structure through synaptic plasticity.

First, we investigate how HOCs affect the development of connectivity in recurrent networks of spiking neurons in the absence of structured external stimuli, where the stochastic activity of each neuron is described by a mutually exciting Hawkes process [67]. We develop a formal analytical framework for the evolution of specific structural motifs in the network based on the second- and third-order moments of spike timing interactions, assuming a slow change of synaptic efficacies following the triplet STDP rule and fast spiking dynamics [52,53]. We demonstrate differences to the classical pair-based STDP rule [50,68] that ignores those HOCs, and compare the relative strength of the emergent structural motifs induced by triplet STDP. Second, we examine the biological conditions which promote the formation of assembly structures of self-connected neurons without externally structured inputs under the triplet STDP rule. We find that this is achieved either by modulating the shape of the STDP function through neuromodulators or the shape of the evoked postsynaptic current (EPSC), and study how the structures formed in these two cases are distinct. Finally, we characterize changes in functional connectivity in terms of graph theoretic measures previously used for describing assembly formation in the zebrafish tectum [25–27].

## Results

We present two main results: first, we derive a formal analytical framework for the evolution of network structures depending on the second- and third-order moments of spike time interactions under the triplet STDP rule; second, we discuss the functional implications of this framework and present the biological conditions which promote the formation of assemblies without external instruction.

### Average synaptic modification due to the interaction of pairs and triplets of spikes in recurrent networks

To study the autonomous emergence of assemblies in a recurrent network from a general form of STDP that includes the contribution of pairs and triplets of spikes to synaptic plasticity, we require a minimal theoretical representation of the network with plastic synapses driven by internal correlations in the spike timing statistics. In our model, structure is given by the connectivity matrix **W** between all excitatory neurons in the network, where the synaptic efficacy element *W_ij_* is different from zero when a connection exists between postsynaptic neuron *i* and presynaptic neuron *j*. The analytical description of the dynamics in recurrent networks can be dauntingly complex. On the one hand, to rigorously analyze the impact of STDP on the formation of functional structures it is indispensable to take into account the precise timing of action potentials or spikes. Therefore, models of neural activity that are based on rates cannot fulfill this criterion [69]. More elaborate models such as Hodgkin-Huxley with multiple ion channels [70] and even the simpler spiking leaky integrate-and-fire (LIF) models are much more accurate in reproducing the spiking dynamics of a population of neurons [71–73]. Although they are computationally tractable, to extract extensive and exact mathematical features from these models remains an elusive task. Under certain conditions of approximately asynchronous firing, the spiking statistics in networks of LIF neuron can be described by a linear theory [29]. Using this approach, here we can make approximations for the spiking dynamics of each individual excitatory cell and treat each pre- and postsynaptic spike trains as if they follow inhomogeneous Poisson statistics [6,44,50,74].

In this model we assume that the probability of each neuron emitting an action potential at a certain time (the ‘intensity’ or mean activity) is proportional to the weighted sum of the preceding activity of all the other cells in the network and a constant, unstructured external input (Fig. 1A). The activity of each neuron in this network is a stochastic process, also referred to as a ‘mutually exciting point process’ or Hawkes process [67]. The availability of an exact expression for spike correlations in this model allows us to develop a precise theory for the synaptic efficacies’ dynamics that are governed by different forms of STDP. To prevent runaway excitation, we also consider that the firing of excitatory neurons is modulated by the activity of a population of inhibitory neurons (Fig. 1A). We do not model plasticity in the connections of inhibitory neurons, since the Hawkes process is formally defined for positive interactions. Nevertheless, we assume that the total inhibitory input to each excitatory neuron is tuned in order to balance the sum of inhibitory efficacies with the sum of the excitatory ones (Methods) [6,75–77].

**Figure 1.**
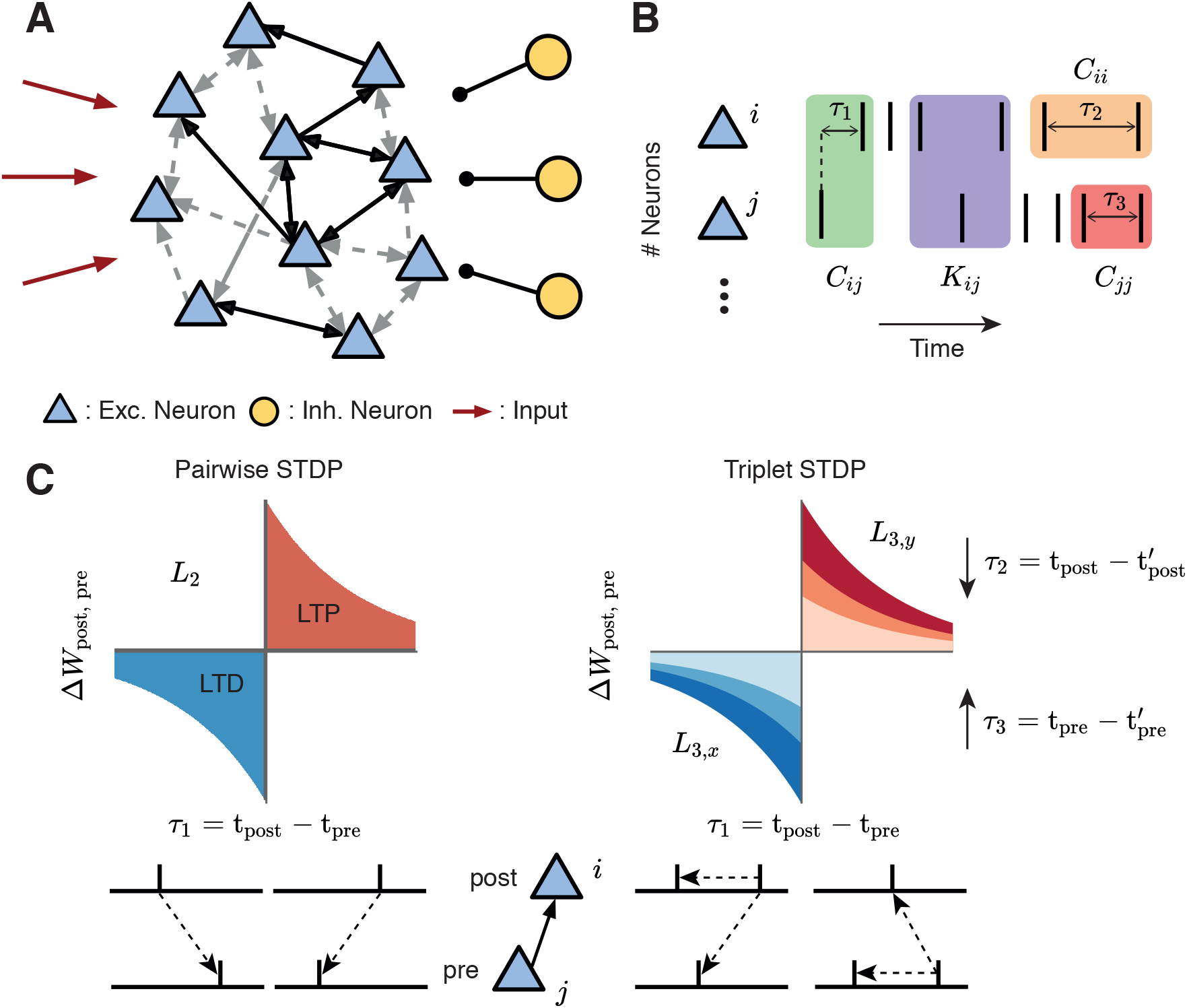
Framework set-up. **A.** A network of excitatory neurons (light blue triangles) fire stochastically, while their activity is driven by unstructured external input (red arrows) and modulated by a population of inhibitory neurons (yellow circles). Excitatory connections among the neurons can be weak (grey dashed arrows) or strong (black solid arrows), unidirectional or bidirectional. **B**. Cumulants of the spike trains (see Eq. 1). The second order cumulants *C_ij_*, *C_ii_* and *C_jj_* are calculated based on the time difference between a pair of spikes (cross-covariance in green; auto-covariances in orange/red). The third-order cumulant *K_ij_* is calculated based on the time differences between three spikes (purple). The spike triplets can be two post- and one presynaptic spikes, or one post- and two presynaptic spikes, depending on the synaptic connection being considered. The time differences are: *τ*_1_ between a presynaptic spike and a postsynaptic spike, *τ*_2_ between different postsynaptic spikes and *τ*_3_ between different presynaptic spikes. **C.** STDP is determined by pairs and triplets of spikes. Left: An example of a classic pair-based STDP rule, with learning window denoted by *L*_2_. Potentiation is triggered by a postsynaptic following a presynaptic spike (*τ*_1_ = t_post_ – t_pre_ > 0), whereas if a presynaptic spike follows a postsynaptic spike (*τ*_1_ = t_post_ – t_pre_ < 0), depression is induced. The total potentiation (depression) is given by the red (blue) area under the curve. Right: Examples of triplet STDP rules denoted by *L*_3,*y*_ and *L*_3,*x*_. Potentiation (red) and depression (blue) are given by triplets of spikes: post-pre-post with a time difference 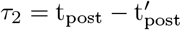, and pre-post-pre with a time difference 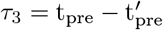, respectively.

Given the connectivity matrix **W** and in the case of a slow learning rate (much slower than the dynamics of neural activity), the rate of change in the strength of synaptic efficacy 〈*Ẇ_ij_*〉 between postsynaptic neuron *i* and presynaptic neuron *j*, can be expressed in terms of the product of the time dependent cumulants of different orders and the STDP function, accordingly (Methods). Specifically, we consider STDP learning rules where plasticity depends on the timing and order of pairs and triplets of spikes, that we call pair-based and triplet STDP. Initially, we make no assumptions about the shape of these learning rules keeping the framework general. The sign and magnitude of the net weight modification depends on the time interval between the firing of the pre- and postsynaptic neurons, and also on the relative spike times of individual pre- and postsynaptic neurons (Fig. 1B). The exact expression for the evolution of the average synaptic efficacy in the recurrent network is

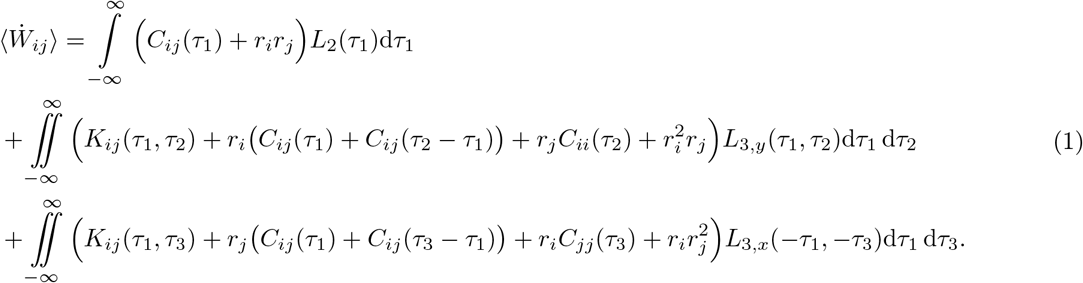

Here *r_i_* and *r_j_* denote the mean firing rates of neuron *i* and *j*, respectively; *C_ij_* is the covariance between neuron *i* and neuron *j*, with *C_ii_* and *C_jj_* being the auto-covariances; and *K_ij_* is the third-order cumulant between neuron *i* and neuron *j*. These quantities represent internal (i.e. not driven by external input) correlations in the network and are calculated as functions of the excitatory postsynaptic current (EPSC), and assumed to be identical for every pair of neurons. Both the covariances *C* and the third-order cumulants *K* are probability densities of pairs and triplets of spikes separated by the given time lapses *τ* accordingly (Fig. 1B). *τ*_1_ is the time difference between a spike emitted by the presynaptic neuron and one from the postsynaptic neuron, whereas *τ*_2_ and *τ*_3_ are the time intervals between different spikes from the postsynaptic neuron and the presynaptic neuron, respectively. The cumulant *K_ij_* is calculated for both ‘post-pre-post’ or ‘pre-post-pre’ spike triplets and therefore depends on combinations of *τ*_1_ and *τ*_2_ or *τ*_3_, according to each case.

The STDP functions that describe how potentiation or depression depend on the spike timing intervals are given by *L*_2_ for pairs of spikes, and *L*_3,*x*_ and *L*_3,*y*_ for triplets of spikes. The sub-indices *x* and *y* correspond to the triplet sets ‘pre-post-pre’ and ‘post-pre-post,’ respectively. While Eq. 1 can be calculated for any shape of the STDP function, an illustrative example for these learning rules, commonly used in other studies based on fits to experimental data [52,53,68], is given in Fig. 1C. The average synaptic efficacy change (Eq. 1) is sufficient to describe the plasticity dynamics when the learning rate is small relative to the spiking dynamics, and noise in the STDP dynamics, arising from random fluctuations, is averaged out. Furthermore, Eq. 1 is combined with heterosynaptic competition [78] to restrict the amount of connections a neuron can make with the rest and prevent the dominance of a few (Methods). For the sake of simplicity, in the next steps we consider that triplets of spikes contribute only to potentiation and thus *L*_3,*y*_(*τ*_1_, *τ*_2_) = *L*_3_(*τ*_1_, *τ*_2_) and *L*_3,*x*_(*τ*_1_, *τ*_3_) = 0, for all *τ*_1_ and *τ*_3_, in agreement with the so-called ‘minimal’ triplet STDP rule [52]. Nevertheless, if spike triplets would also be taken into account for depression, the derivation would be identical, with the corresponding modification to the variables involved. After some calculations, we can rewrite Eq. (1) in the Fourier domain as

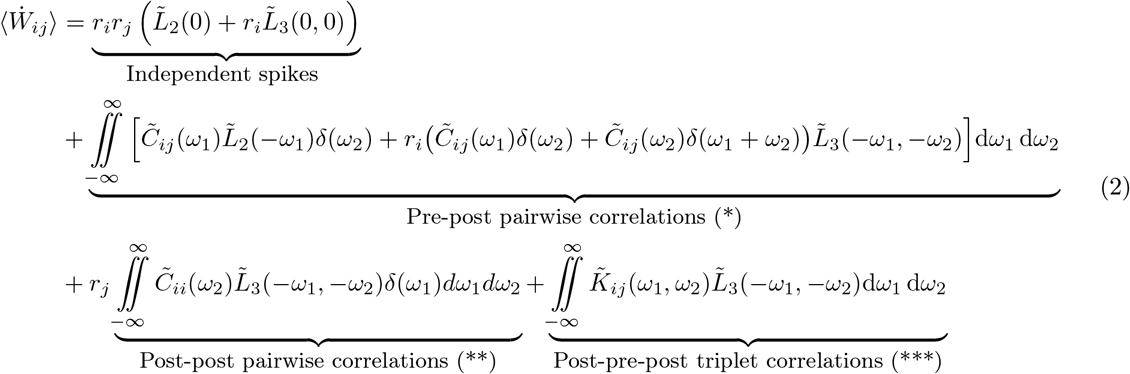

where we use the notation 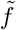 for the Fourier transform of a function *f* and *δ* is the Dirac delta function. It should be noted that Eq. 2 is not the Fourier transform of Eq. 1 but rather an equivalent expression of the latter. This comes about because we can express the integral of the product of two functions as the convolution of the Fourier transform of those functions, evaluated at zero. This formulation of the previous equation allows us to clearly break down the contribution of spike interactions of different orders to the average synaptic efficacy in the recurrent network. The first term of Eq. 2 considers the change in synaptic efficacy that is obtained from independent spiking and thus depends on the first-order moments (the mean firing rates) of the activity of both the pre- and postsynaptic neurons, *r_j_* and *r_i_*, respectively. As firing rates increase, ‘chance’ contributions to plasticity can occur. The second and third term account for the probability of observing changes to the mean synaptic efficacy due to pairwise correlations in the pre- and postsynaptic neurons. *C_ij_* refers to the family of probabilities that generate pairwise correlations (second-order moments) between neurons *i* and *j*, depending on spikes of other neurons in the network (Fig. 1B, green). Accordingly, *C_ii_* includes the family of probabilities that generate pairwise auto-correlations in the same neuron *i* due to the spiking activity of all other neurons in the network (Fig. 1B, orange). Therefore, the second (*) and third (**) terms describe the total contribution of correlated spike pairs to plasticity through the pair-based STDP rule *L*_2_ (Fig. 1C, left) and the triplet STDP rule *L*_3_ (Fig. 1C, right). In the case of the latter, the first-order moment of the uncorrelated single postsynaptic neuron’s spikes, *r_i_*, is also included in the second term (*) and the first-order moment of the uncorrelated single presynaptic neuron’s spikes, *r_j_*, in the third term (**). The fourth term (***) describes the total contribution of correlated spike triplets (third-order moments) to plasticity. Thus, *K_ij_* includes the family of probabilities for third-order correlations, where the relative spike timing interacts with the triplet STDP learning window *L*_3_ to induce plasticity (Fig. 1B, blue and Fig. 1C, right). In conclusion, we have derived an exact analytical expression for the average change in synaptic efficacy due to firing rates, pairwise and triplet correlations under a general STDP rule that includes pairwise and triplet spike interactions.

### Novel branching structures emerge under triplet STDP compared to pair-based STDP

The expansion involving second- and third-order cumulants in the equation for the average weight dynamics (Eq. 2) can be used to determine the evolution of ‘wiring’ or ‘structural’ motifs – specific patterns of connectivity among groups of neurons – that arise from implementing the pair-based and triplet STDP rules (examples given in Fig. 3A,C). These motifs are over- or under-represented in neural circuits compared to Erdös-Rényi (random) networks [5,16,51,79], and their predominance can be linked to shared stimulus preference and elevated levels of activity [5,13,14,80]. The influence or ‘drive’ of each particular motif on the structure that emerges in the network is determined by the ‘motif coefficient’ of each particular motif. Then, to calculate each term in Eq. 2 we break down the moments *C_ij_, C_ii_* and *K_ij_* into expressions that include the contribution of every spike propagated in the network through existing synaptic connections (Methods). These expressions consist of products of the corresponding synaptic efficacies from the connectivity matrix **W** and the motif coefficient functions *M*, which depend on the EPSC function, *E*, and the STDP learning rules, *L*_2_ and *L*_3_. The probability that neurons *i* and *j* jointly fire a spike is transiently modulated whenever a neuron anywhere in the network produces a spike. Each term in the pairwise cross-covariance from Eq. 2

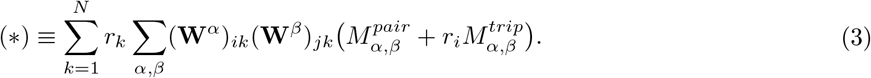

quantifies how this modulation results in a change in the connectivity matrix **W**. The expression consists of two sums to provide an intuitive description of the contribution of the pairwise cross-correlation *C_ij_* between neurons *i* and *j* to plasticity of the connection *W_ij_*. The first sum takes into account all spiking neurons in the network, while the second sum takes into account all possible ‘paths’ by which spikes originating from a ‘source’ neuron *k* affect the pairwise correlation *C_ij_*. Specifically, *α* and *β* constitute the ‘path lengths’ of synapses from source neuron *k* to the postsynaptic neuron *i* and the presynaptic neuron *j*, respectively (Fig. 2A; see also [6]).

**Figure 2.**
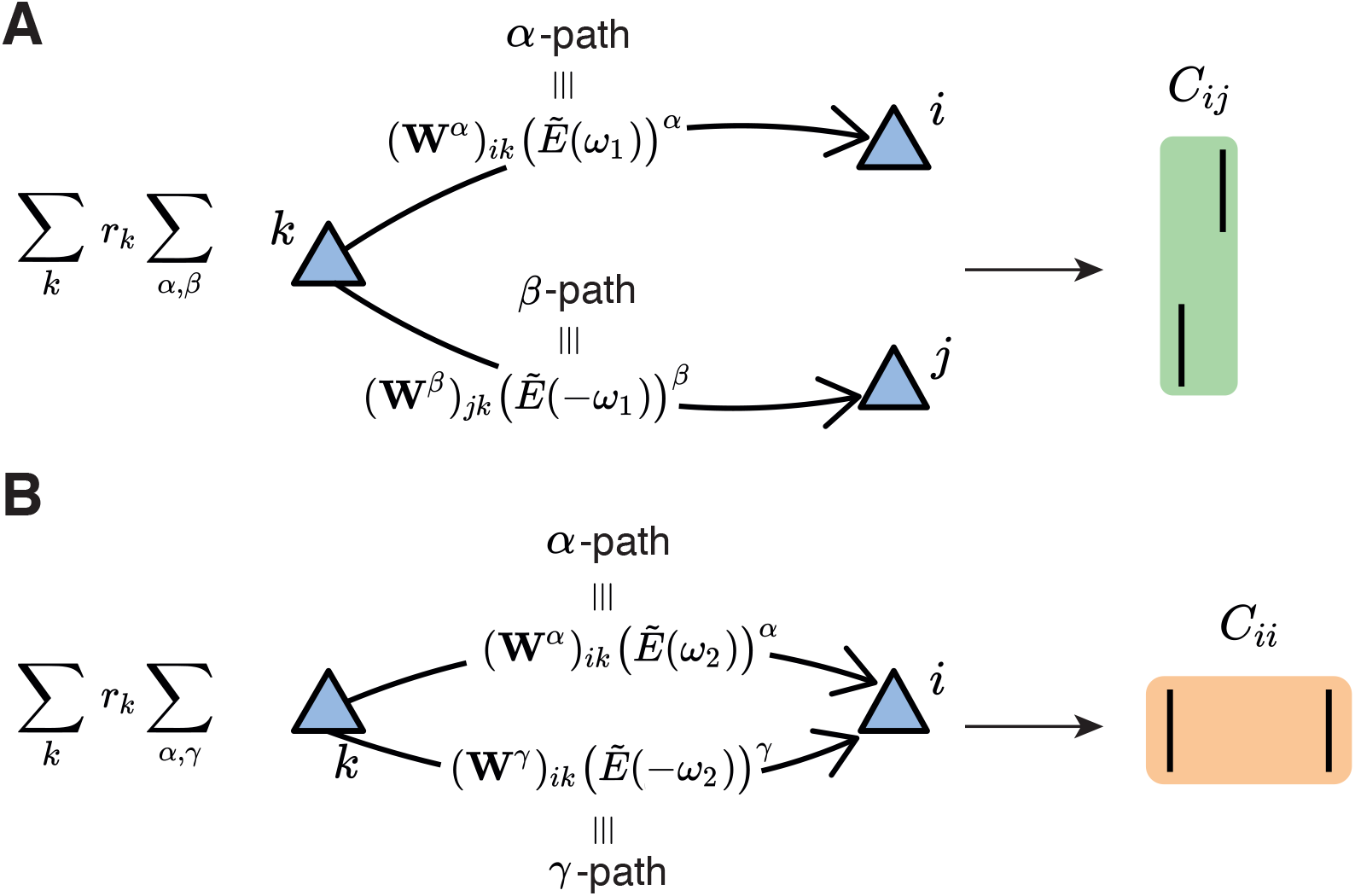
Second-order cumulant contributions to plasticity. **A.** The cross-covariance *C_ij_* between the presynaptic neuron *j* and the postsynaptic neuron *i* is obtained by summing over all the possible *α*- and *β*-paths from every possible source neuron *k* in the network. Each path is calculated via the corresponding weights in the connectivity matrix and the EPSC function (see Eq. 3). **B**. Same as **A** but for the auto-covariance *C_ii_* of the postsynaptic neuron *i* (see Eq. 4). In this case, *γ* is the second index to sum over the path from the source neuron *k* to the postsynaptic neuron *i*. It should be noted that the main difference between the *α*- and *γ*-path is given by the time dependence of the EPSC function, here represented in the Fourier domain for convenience.

**Figure 3.**
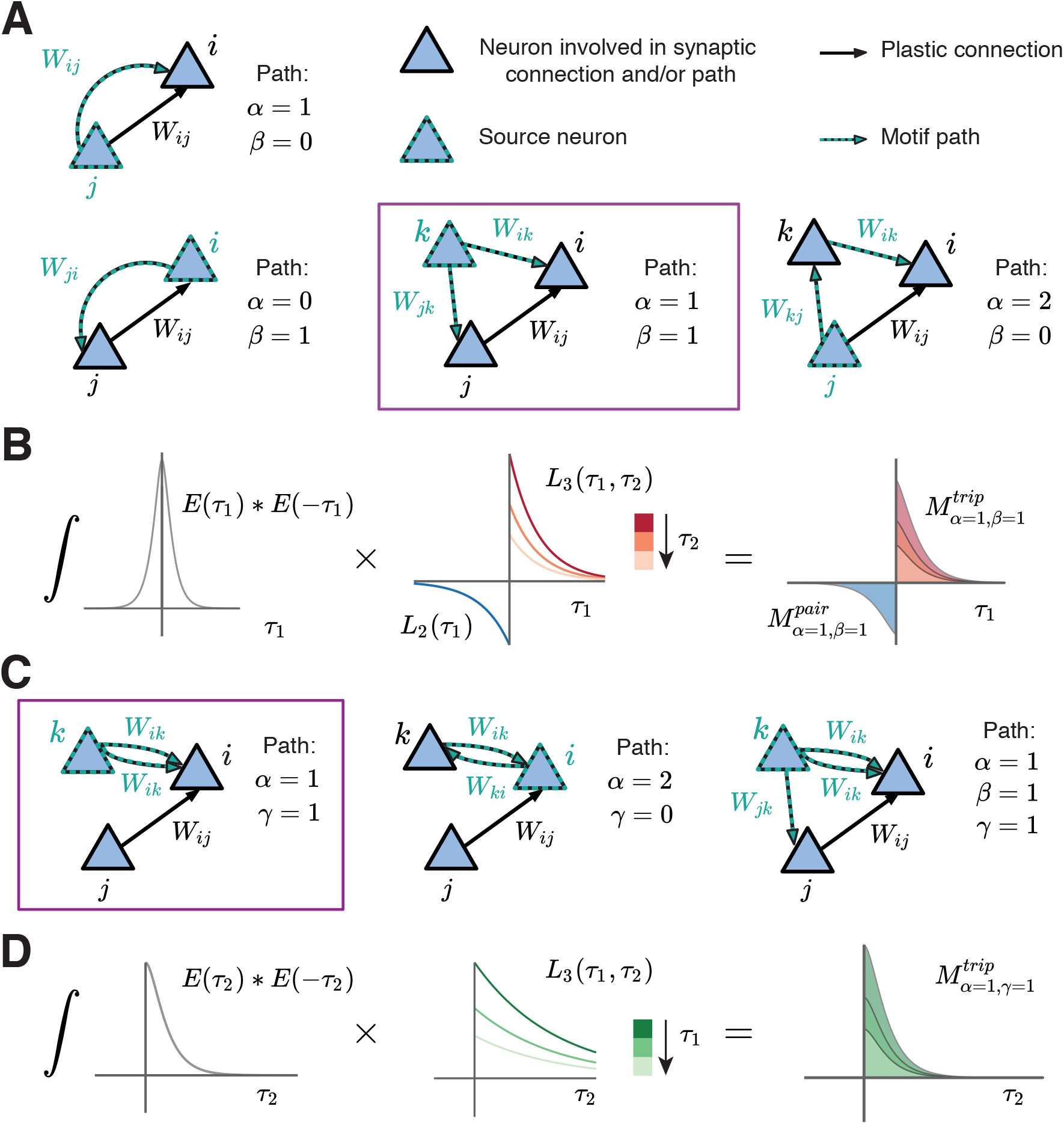
Novel motifs in the network under triplet STDP. **A.** Examples of motif structures common for both the pair-based and triplet STDP framework. Here *α* and *β* constitute the path lengths of synapses from the source neuron to the postsynaptic neuron *i* and the presynaptic neuron *j*. *α* = 1, *β* = 0: Presynaptic neuron *j* projects to the postsynaptic neuron *i*. *α* = 0, *β* =1: Postsynaptic neuron *i* projects to the presynaptic neuron *j. α* = 1, *β* = 1: Common input from source neuron *k* to presynaptic neuron *j* and postsynaptic neuron *i. α* = 2, *β* = 0: Presynaptic neuron *j* projects to the postsynaptic neuron *j* through another neuron *k* in the network. **B.** Illustration of the calculation of the common input motif with *α* = 1 and *β* =1 framed in purple in A (additional terms not shown). The motif coefficients *M*_*α*=1, *β*=1_ (right) are calculated as the total area under the curve resulting from the product of the convolution of the EPSC function *E* (left) and the STDP functions (pair-based *L*_2_ and triplet *L*_3_, middle). **C.** Examples of motif structures found only in the triplet STDP framework, where *γ* denotes the time-delayed path length from the source neuron to the postsynaptic neuron *i*. *α* = 1, *γ* =1: Source neuron *k* projects twice to postsynaptic neuron *i* with a different time delay. *α* = 2, *γ* = 0: Feedback loop through another neuron *k* in the network (source and projecting neuron are the postsynaptic neuron *i*). *α* = 1, *β* = 1, *γ* =1: Source neuron *k* projects to the presynaptic neuron *j* and postsynaptic neuron *i* via all the three possible paths. **D.** Illustration of the calculation of the motif with *α* = 1 and *γ* = 1 for the triplet STDP rule framed in purple in **C**, compare to **B**.

The contribution of the pair-based STDP rule includes the motif coefficient functions, 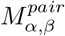, which are calculated in the Fourier domain (Eq. 31 in Methods). The pairwise correlations between *i* and *j* also contribute to plasticity of *W_ij_* based on the triplet STDP rule through the motif coefficient functions 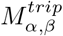 (Eq. 32 in Methods). Examples of some motifs common for both the pair-based and the triplet STDP rule are provided in Figure 3A. Their contribution to plasticity through the EPSC function *E* and the STDP rules *L*_2_ and *L*_3_ is illustrated in Figure 3B.

In addition to the *α* and *β* path lengths, to derive the contribution of the triplet STDP rule to the average change in synaptic efficacy, we also introduced the *γ*-path. *γ* is the synapse path length from the source neuron *k* to the postsynaptic neuron *i*, including a time delay relative to the *α* path from *k* to *i*, to account for the second postsynaptic spike of the triplet (Fig. 3C). Thus, for the auto-covariance term in Eq. 2, we obtain (Fig. 2B)

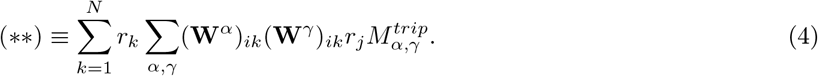

where the motif coefficient function involving the triplet STDP rule is given in the Methods (Eq. 33).

For third-order interactions, however, it is possible that the paths by which spikes are propagated branch out from a neuron other than the source neuron. Therefore, the cumulant *K_ij_* (Eq. 2) is broken down into four sums:

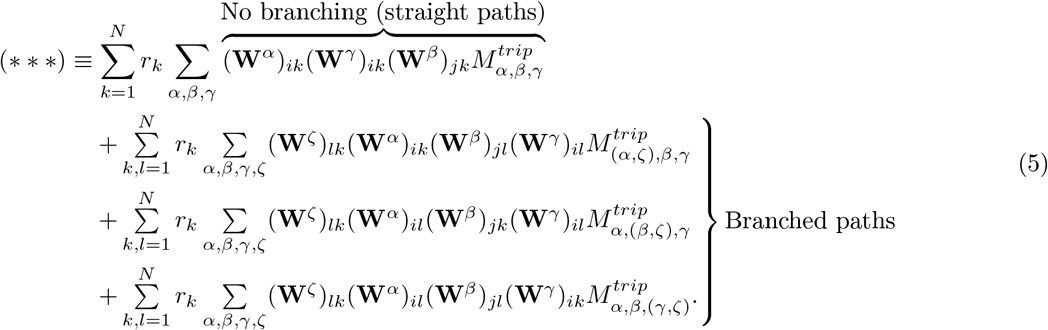

The first term in Eq. 5 sums over the paths to the presynaptic neuron *j* and postsynaptic neuron *i* from a source neuron *k* in the network that do not branch out. In other words, it considers that the ‘distance’ to each respective spike of the triplet is given by *α*, *β* and *γ* (Fig. 4A). The remaining terms include the sum over possible branches in the network ‘tree’: *ζ* ≥ 1 is the synapse path length from the source neuron *k* to the neuron *I* that is the branching point (Fig. 4B-D). It should be noted that this is only possible for motifs of order higher than three, since at least one synapse must be taken into account before the splitting of the path. The corresponding motif coefficients for the ‘straight’ triplet motif (Fig. 4A, see Eq. 34), and for the ‘branching’ motifs (Fig. 4B-D, see Eqs. 35, 36 and 37) are provided in the Methods.

**Figure 4.**
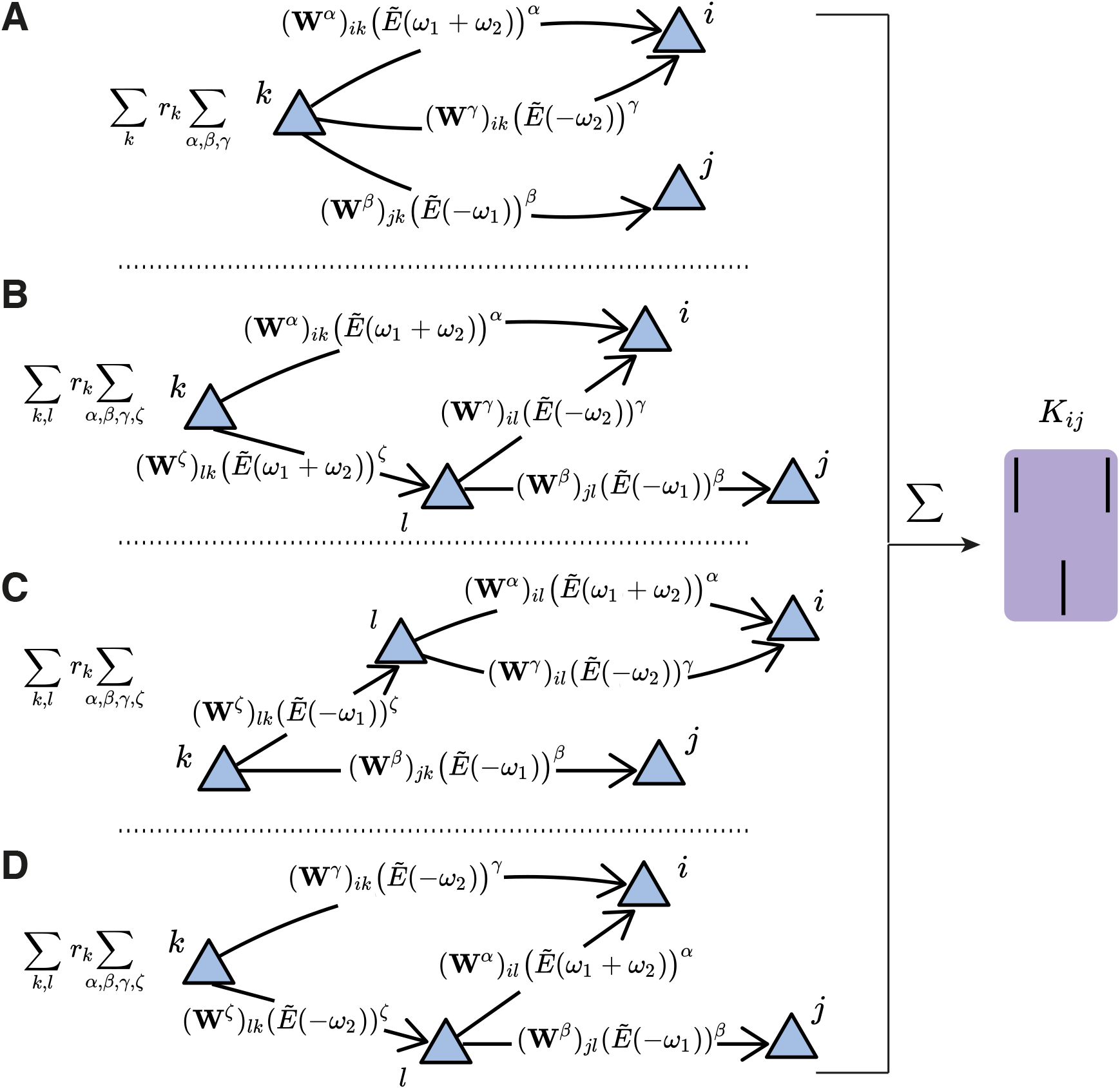
Third-order cumulant contributions to plasticity can be broken down into four terms. **A.** The first term contains all the *α*-, *β*- and *γ*-paths originating from the source neuron *k* to the spiking neurons *i* and *j*. **B-D.** The other terms take into account the possibility of an intermediate neuron *l* that acts as a new source neuron for two of the paths. These are referred to as ‘branching paths’, and the path length from the source neuron *k* to the intermediate neuron *l* is denoted with *ζ*. The branching describes the individual terms in Eq. 5.

This analysis reveals that spike triplet interactions in the STDP rule can promote particular connectivity structures that are not possible with pairwise interactions [6] (Fig. 3C). These include motifs which directly exclude the presynaptic neuron *j* but can still impact the synaptic weight, *W_ij_* (Fig. 3C, left and middle). This can be achieved, for example, through an additional neuron *k* that does not directly affect the weight *W_ij_* but projects to the postsynaptic neuron *i* though the synaptic weight *W_ik_* (Fig. 3C, left and middle). For example, in the case when *α* = 2 and *γ* = 0 (Fig. 3C, middle), the postsynaptic neuron *i* is both the source neuron and the neuron involved in the path with the additional neuron *k*. We call this path involving the synaptic efficacies *W_ik_* and *W_ki_* a ‘loop’. Motifs with these characteristics act as a nonlinear filter for the postsynaptic firing rate (Eq. 18), and promote the formation of connections between clusters of neurons, and therefore assemblies. These loops involving a different neuron in addition to the pre- and postsynaptic neuron involved in the weight *W_ij_* are a unique feature of incorporating spike triplets in the STDP rule and are distinct from the bidirectional connections arising from motifs containing the pre- and postsynaptic neuron. In the latter case, promoting the motifs *α* = 1, *β* = 0 and *α* = 0, *β* = 1 (Fig. 3A, left) supports the formation of a bidirectional connection on that synapse (*W_ij_* and *W_ji_*; see next section for details). However, since the source neuron is different for each motif in this case, the resulting bidirectional connection is not equivalent to a loop. Taken together, our motif expansion framework reveals novel motif structures under the triplet STDP rule that enable the formation of assemblies, in comparison to the bidirectional connections under the pair-based STDP rule.

### Modulation of the triplet STDP rule promotes the autonomous emergence of assemblies

So far, we considered general STDP rules that depend on the precise timing between pairs and triplets of spikes, without taking into account the exact dependence of potentiation or depression on these spikes. To further study the complex relationship between plasticity and network correlations, we considered a particular biologically identified STDP rule that relies on third-order interactions (Methods; Fig. 1C). This rule has an asymmetric shape around the time lag of 0 (where pre- and postsynaptic spikes are coincident), similar to the classical pair-based STDP rule [68]. However, while synaptic depression is induced by the relative timing of pairs of presynaptic and postsynaptic spikes, the triplet STDP model uses sets of three spikes to induce potentiation: the amount depends on the timing between pre- and postsynaptic spike pairs and in addition, on the timing between the current and the previous postsynaptic spike. This *minimal* model *successfully* captures experimental data where the pairing frequency of pre- and postsynaptic spikes was varied equally well compared to a full model that also used triplets of spikes for depression [52].

Implementations of classical Hebbian learning, such as STDP, use joint pre- and postsynaptic activity to induce potentiation and depression, while neglecting other potential factors such as heterosynaptic plasticity [81], or the location of synaptic inputs on the dendritic tree [82]. However, recent experimental studies have highlighted an important role of neuromodulators in regulating plasticity across the brain [83–86], as they convey information about novelty or reward. Indeed, neuromodulators such as dopamine, acetylcholine and noradrenaline, but also brain-derived neurotrophic factor (BDNF) and gamma-aminobutyric acid (GABA), can predominantly act via two mechanisms: by reshaping the learning window for STDP or by regulating the neuronal activity at the level of synaptic transmission [84,86]. Therefore, we next investigated how neuromodulation of synaptic plasticity affects recurrently connected networks considering that pairwise and triplet spike interactions determine plasticity. We assume that the shape of the STDP function can be modulated via the modulation parameter *η*_−_ which preserves the overall level of depression by trading off the depression learning rate *A*_−_ and the depression time constant *τ*_−_ (Methods; Fig. 5A). Such a modification of the learning rule has been observed in the lateral amygdala due to the action of dopamine via D2 receptors [85,87], or in rat visual cortex slices with the activation of both the noradrenaline pathway through *β*-adrenergic receptors and the acetylcholine pathway through *M*_1_-muscarinic receptors [84,86,88]. A similar modulation parameter could similarly be included for potentiation.

**Figure 5.**
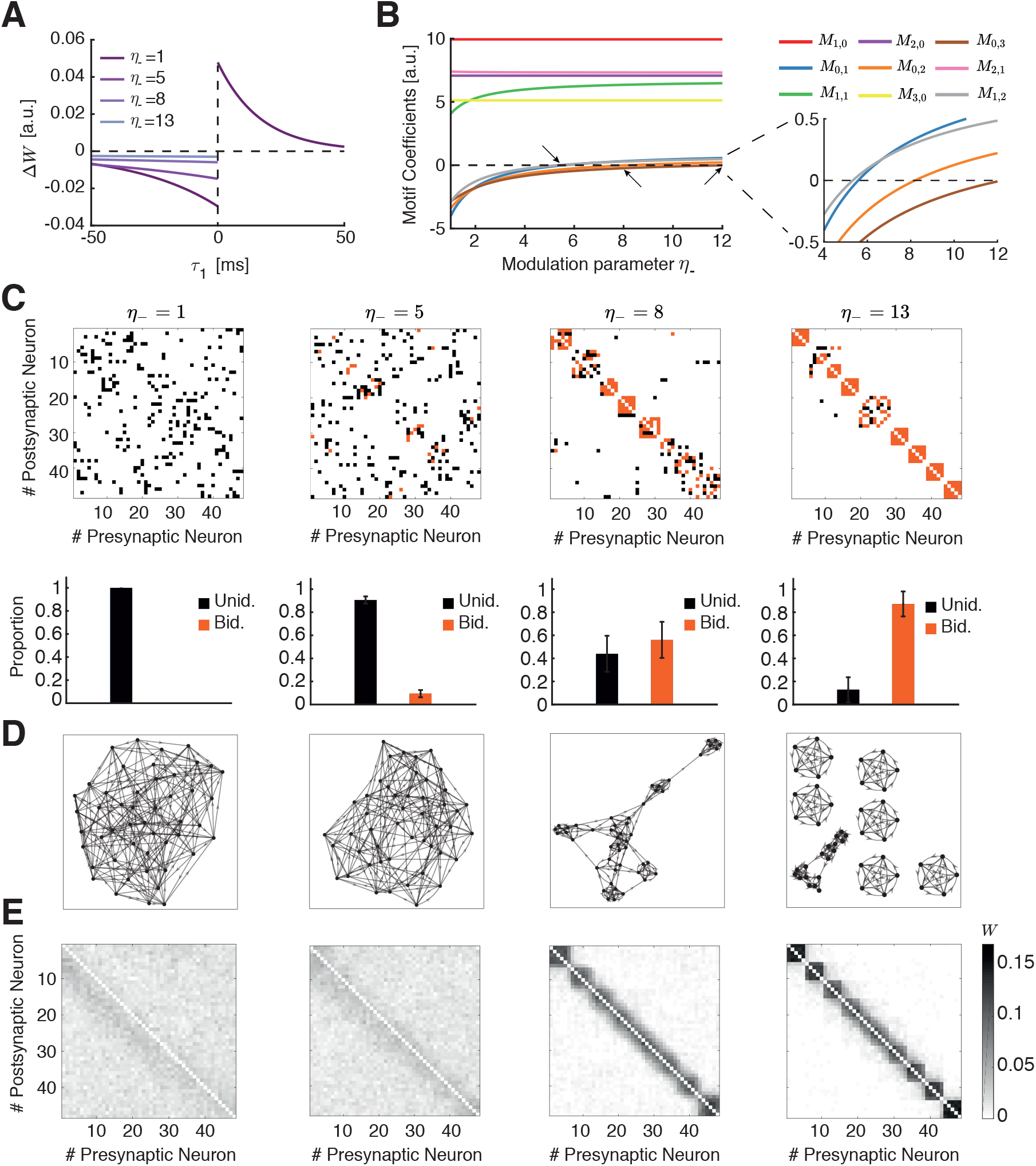
Spontaneous emergence of assemblies via modulation of the triplet STDP rule. **A.** The shape of the STDP function changes as a function of the modulation parameter *η*_−_. **B**. Motif coefficients as the modulation parameter *η_−_* increases. Points of interest given by the crossovers of the strength of particular motifs are indicated by a small arrow. Inset: Amplified scale around zero. Motif coefficients including *γ* paths are not illustrated, since they are always constant and positive in *η*_−_ and do not provide a meaningful comparison to the other motifs. **C.** Top: Examples of connectivity matrices obtained with different values of *η*_−_. Unidirectional connections are shown in black, bidirectional connections in orange. Bottom: Proportion of unidirectional and bidirectional connections over total connections, for 100 trials with different initial synaptic efficacies as a function of *η*_−_. **D.** Graphs of the connectivity matrices in **C. E.** Averaged connectivity matrices over 100 trials. Note that the tighter clusters emerging near the edges of the matrices are the result of the clustering algorithm but do not affect quantification of connectivity (see Methods).

To determine contributions to plasticity arising due to internal network correlations and not just differences in neuronal firing rates [5], we consider the case in which the plasticity rule is balanced, such that 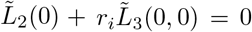. We use this condition to calculate all motif coefficients, *M_α,β_*, that arise from pairwise and triplet correlations (Eqs. 49–58 in Methods). We consider only motifs up to third-order in the evolution of the weights (Eq. 2) and thus no longer include the branched path motifs of Eq. 5 as they are higher-than-third-order and have weaker influence on weight evolution (Fig. 4B-D).

To systematically study how the dependence of these motif coefficients on the shape of the STDP rule affects connectivity structure in the network, we visualized the connectivity matrices obtained by integrating Eq. 2 numerically, using experimentally-fitted parameters for the triplet STDP rule and the EPSC function (Table 1). Specifically, we investigated the emergence of global network structures, or assemblies, as a function of the modulation parameter *η*_−_. A key requirement for the emergence of assemblies is the formation of bidirectional or reciprocal connections among groups of neurons. We compare the reciprocal connections of the first-order motif contributions to gain intuition:

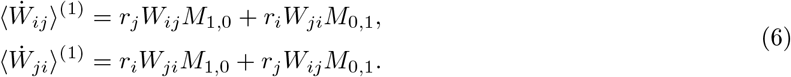

**Table 1.**
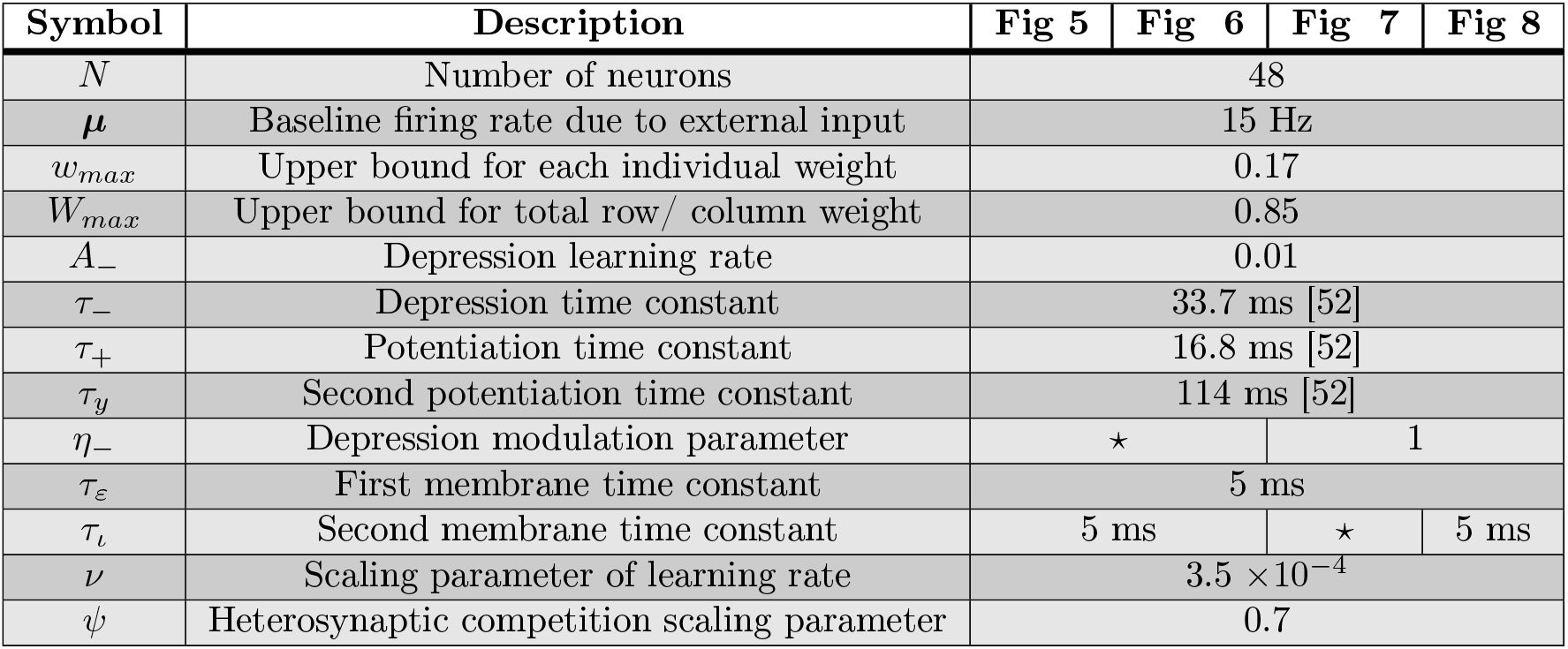
Parameters values for figures. ⋆ denotes that values are provided in the figures.

Since in the triplet STDP rule *M*_1,0_ > 0 (Fig. 5B, red), the two bidirectional connections compete if *M*_0,1_ < 0, and potentiate if *M*_0,1_ > 0. Therefore, the sign of the motif coefficient *M*_0,1_, which depends on the corresponding pairwise and triplet correlation contributions, determines the formation of bidirectional connections. Indeed, increasing *η*_−_ supports the formation of bidirectional connections (Fig. 5C) as the motif coefficient *M*_0,1_ changes sign (Fig. 5B, blue, see inset). In contrast, as previously shown, the classical pair-based STDP rule is unable to support the formation of assemblies and bidirectional connections due to its asymmetric shape [89,90], although under certain conditions (dominant potentiation) it can promote bidirectional connections as local two-synapse connectivity motifs that ignore network interactions [91]. Under the asymmetric pair-based STDP rule, *M*_1,0_ > and *M*_0,1_ < 0 result in competition between the two reciprocal connections. To autonomously generate self-connected assemblies without structured network input requires a symmetric pair-based STDP rule (which is not biologically motivated) and a sufficiently large synaptic latency [6]. In this case, the prominence of the common input motif driven by the *M*_1,1_ motif coefficient, over all other motif coefficients in the network supports assembly formation [6].

Under the triplet STDP rule, small increases in *η*_−_ increase the motif coefficient *M*_1,1_, resulting in the formation of bidirectional connections and assemblies, similarly to the symmetric pair-based STDP rule. However, despite its asymmetric shape, the triplet STDP rule can robustly generate bidirectional connections and assemblies even when the *M*_1,1_ motif coefficient has already statured and other motif coefficients dominate (Fig. 5C-E), upon further increases in *η*_−_. This is because higher-order motif contributions also contribute to the formation of bidirectional connections and assemblies. To understand this, we consider the motif contributions of feedforward motif coefficients – the motifs for which the *α*-path is longer than the *β*-path – and reciprocal motif coefficients, where the *β*-path is longer than the *α*-path. Given the asymmetry of triplet STDP, the feedforward motif coefficients are the strongest. The reciprocal motifs, *M*_0,1_, *M*_1,2_, *M*_0,2_ and *M*_0,3_ play an important role in the formation of bidirectional connections as they change sign from negative to positive with increasing *η*_−_ (Fig. 5B). A positive contribution from all motifs supports the robust formation of bidirectional connections in the network as the competition between reciprocal connections decreases. Together with the strong common input motif driven by the *M*_1,1_ motif coefficient, this leads to the emergence of assemblies. In this scenario, *η*_−_ controls the competition between feedforward (*W_ij_*) and reciprocal connections (*W_ij_*), with large *η*-enabling the potentiation of both. This is not possible under the classical asymmetric pair-based STDP rule as previously discussed.

In summary, we find that self-connected assemblies can be formed even when the common input motif is not dominant when considering the triplet STDP rule. This occurs despite the asymmetric shape of the triplet STDP rule. Importantly, the dependence of assembly formation on the specific form of the STDP window points towards an important role of neuromodulatory signals on formation of intrinsically generated assemblies.

### Characterizing emergent assembly structures

To determine the conditions on the learning rule and EPSC properties for the emergence of self-connected assemblies, it is convenient to represent the functional organization of the network given a connectivity matrix as a mathematical graph. In our context, graphs are composed of a set of nodes or neurons with pairs of them joined by edges or synaptic efficacies. The resulting graphs can be described by standard metrics, whose dependence on the learning rule and the EPSC function might yield insight into the emergent structures from different forms of plasticity during circuit organization driven by spontaneous activity in experimental systems like the zebrafish tectum or the mammalian cortex [25,92]. We focused on common quantities for describing graph structure, including the clustering coefficient, the global efficiency and the modularity [93,94].

The clustering coefficient quantifies the existence of densely interconnected groups of nodes in the graph [95]. It represents a measure of segregation, based on counting the number of connection triangles around a node (Methods). In this manner, it reflects the prevalence of clustered connectivity around individual nodes by calculating the fraction of neighbors of that particular node that are also neighbors of each other. As a result, the mean clustering coefficient of a network determines the prevalence of three-neuron-clusters in the network architecture. We find that as the modulation parameter *η*-increases, the mean clustering coefficient also increases until it reaches a plateau (Fig. 6A). Ensuring that the motif coefficients *M*_0,1_ and *M*_1,2_ are positive is sufficient for the formation of clusters beyond the critical value of *η*_−_ ≈ 5 (Fig. 5B and C), where the clustering coefficient begins to increase Fig. 6A). The value of *η*_−_ at which the clustering coefficient saturates corresponds to the emergence of more robust assemblies where all the motif coefficients are positive (Fig. 5B and C). Although strong bidirectional connections are localized within clusters, connections from one cluster to some others still exist globally. This is different to the clustering enabled by strong symmetric interactions in which the motif *M*_1,1_ dominates, considered previously by a symmetric pair-based STDP rule [6], where the clusters would be unconnected (i.e. isolated from each other) and the clustering coefficient would be much higher.

**Figure 6.**
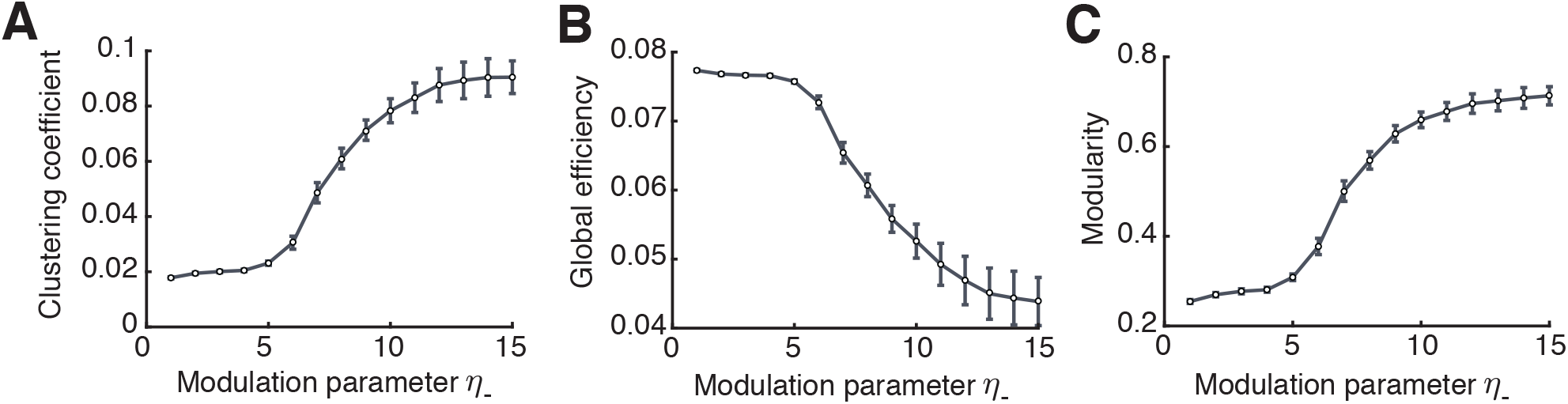
Graph measures of the stable configuration of the networks. **A.** Mean clustering coefficient versus the modulation parameter *η*_−_. **B.** Mean global efficiency versus the modulation parameter *η*_−_. **C.** Modularity versus the modulation parameter *η*_−_. All results calculated from the 100 trials.

Complementary to the clustering coefficient, the global efficiency is a measure of functional integration, which determines how easily nodes can communicate between each other through sequences of edges [96]. Consequently, the lengths of the paths estimate the potential for the flow of information between nodes, with shorter paths denoting stronger capacity for integration. Then, global efficiency is defined as the average inverse shortest path length of the network (Methods). In comparison to the clustering coefficient, this quantity initially remains approximately constant and then decreases until the point at which strong assemblies emerge autonomously since network structure no longer varies with the parameter *η*_−_ (Fig. 6B). We find that as for the clustering coefficient, the value of *η*_−_ for which the motif coefficients *M*_0,1_ and *M*_1,2_ become positive (*η*_−_ ≈ 5) constitutes a landmark for the formation of assemblies, after which global efficiency significantly decreases.

Finally, modularity is a graph-theoretic measure that describes how strongly a network can be divided into modules, by comparing the relative strengths of connections within and outside modules to the case when the network had weights chosen randomly [94,97,98]. Recently, it was shown that even in models with rate-based dynamics, increasing modularity amplifies the recurrent excitation within assemblies evoking spontaneous activation [46]. Thus, it becomes a relevant quantity for characterizing the functional organization of the network. With increasing η_−_, modularity increases until strong assemblies are formed in a similar fashion as the clustering coefficient (Fig. 6C).

### The triplet STDP rule and the EPSC together modulate network structure

Regulating the synaptic transmission of action potentials between neurons through the shape of the EPSC function also provides a way to modulate the overall effect of motifs on network structure. In this manner, the strength of internal correlations can be changed independently of the STDP functions *L*_2_ and *L*_3_. We investigated how the rise of the EPSC function modulated by the parameter *τ_ι_*, which represents the delay of the spike transmission in the synapse (Fig. 7A), shapes these motif coefficients (Methods; Fig. 7B).

**Figure 7.**
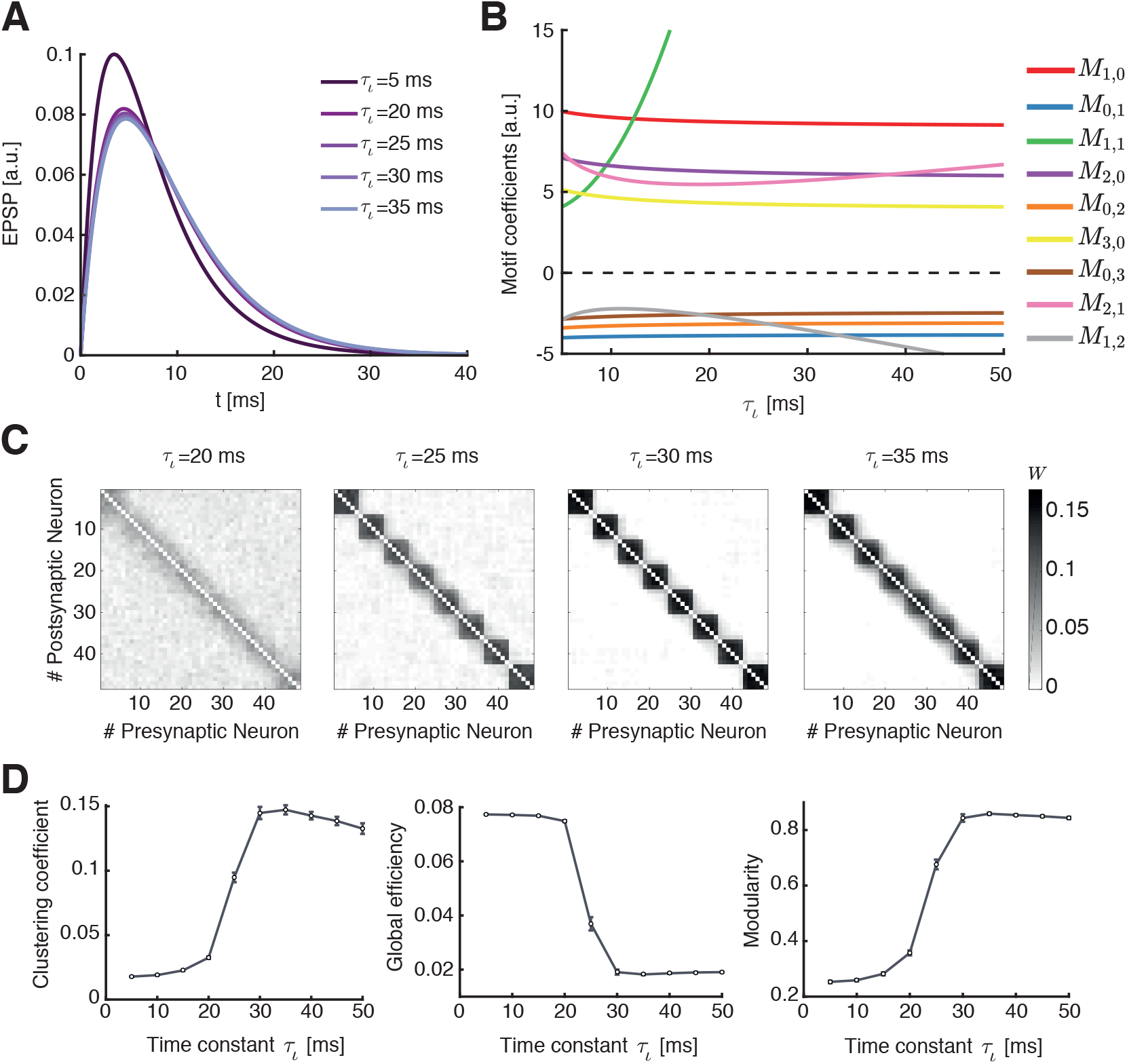
Autonomous emergence of assemblies due to the modulation of synaptic spike conductance. **A.** Varying the time constant *τ_ι_* changes the shape of the time dependency of the EPSC function, shifting its maximum point by a few millisecons. **B.** Relative value of the motif coefficients as the second time constant of the EPSC function *τ_ι_* is increased. It can be easily seen that the common input motif becomes rapidly dominant, but also there exist crossovers of the motif coefficient *M*_1,2_ with the strength of the feedback motifs *M*_0,1_, *M*_0,2_ and *M*_0,3_. **C.** Averaged connectivity matrices for a network of 48 neurons, for 100 trials and various values of the time constant *τ_ι_*. From left to right: *τ_ι_* = 20 ms; *τ_ι_* = 25 ms; *τ_ι_* = 30 ms; *τ_ι_* = 35 ms. The refinement of the self-connected assemblies seems to be symmetric for all clusters, in comparison to the ones obtained with the modulation of STDP. **D.** Clustering coefficient, global efficiency and modularity versus the time constant *τ_ι_*.

The parameter *τ_ι_* has a prominent impact on plasticity in the network. Even small shifts in the peak of the EPSC function by a few milliseconds have a strong impact on the internal correlations calculated in our framework, as reflected in the values of the motif coefficients (Fig. 7A and B). It is evident that the motif coefficient of the common input motif *M*_1,1_ abruptly assumes dominance over all others as *τ_ι_* increases. Thus, assemblies can emerge robustly even in the absence of structured external input or without a symmetric pair-based STDP rule [6]. However, we observed that the reciprocal motif coefficients *M*_0,1_, *M*_1,2_, *M*_0,2_ and *M*_0,3_ remain negative for all values of *τ_ι_*, in contrast to when we modulated the STDP learning rule (Fig. 5B). This tells us that assemblies in the network are spontaneously formed in a different fashion than when modulating the STDP rule. In fact, assemblies emerge for minor modulations in *τ_ι_* (Fig. 7C).

These differences in assembly formation become apparent when we consider the mean clustering coefficient, the global efficiency and the modularity as functions of *τ_ι_* (Fig. 7D): the three measures reflect the connectivity matrices as *M*_1,2_ crosses the motif coefficients *M*_0,1_, *M*_0,2_ and *M*_0,3_, in the case when *M*_1,1_ is already large. When the motif coefficient *M*_1,2_ becomes more negative than *M*_0,3_ (*τ_ι_* ≈ 20 ms) bidirectional connections are strongly promoted and assemblies robustly form. Even for *τ_ι_* > 20 ms, where the EPSC function does not change significantly (Fig. 7A), one sees noticeable changes in the ‘tightness’ of the assemblies as observed in the averaged connectivity matrices (Fig. 7C). Interestingly, as *M*_1,2_ decreases below *M*_0,2_ (*τ_ι_* ≈ 25 ms), the value of the clustering coefficient (≈ 0.1) and the modularity (≈ 0.7) correspond to the values where the clustering coefficient, the modularity, and the global efficiency saturate when modulating the STDP function (compare Fig. 6 and Fig. 7D). This means that the network structure is very similar (compare Fig. 5E, right, with Fig. 7C, second from left). Nevertheless, further increasing *τ_ι_* leads to more refined assemblies (Fig. 7C, third from left) when *M*_1,2_ < *M*_0,2_. However, for *τ_ι_* ≳ 35 ms where *M*_1,2_ < *M*_0,1_, the clustering coefficient slightly decreases (Fig. 7D, left) suggesting the existence of optimal regions in the parameter space of *τ_ι_* to obtain the ‘tightest’ assemblies.

Taken together, our analytical framework enables us to interpret changes in the motif coefficients as changes in the connectivity structure in terms of the formation of self-organized assemblies. Modifying either the shape of the learning rule, or the shape of the EPSC function, can achieve this, however, with different consequences on the nature of the formed structures as demonstrated by the graph theoretic measures.

### Comparison with assemblies generated via external correlated input

Until now, we sought to understand the mechanisms that contribute to the autonomous emergence of assemblies in neural circuits without any structured external input. Yet, the training of assemblies and plasticity of recurrent connections has been more frequently studied when these networks are driven by structured external input, both in simulations [47,89] and analytically [42–45,49]. Significant experimental evidence also exists for the emergence of functional connectivity underlying feature selectivity in the visual cortex around the time of eye opening, which is presumably influenced by structured visual input through the open eyes [14]. Therefore, we wanted to compare the formation of assemblies without structured external input under the triplet STDP rule to that with structured external input. To investigate spatiotemporal input patterns in our framework, we studied the overall mean impact of an external pairwise correlated input. This was implemented by assuming that the driving signal, which could for instance represent retinal input in the optic tectum or visual cortex, is correlated for a pair of neurons in the network, so that the structure of the input is represented as common input to that particular pair of neurons.

We write the pairwise covariance as a sum of the internal correlation and a novel term that conveys the external structured activity as common input [40]:

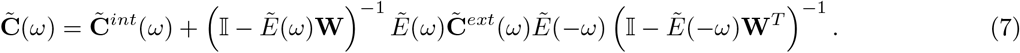

Here, **C**^*int*^ denotes the covariance matrix (see Eq. 24) and **C**^*ext*^ is the covariance matrix of the external input. We model the input signal as a correlated pattern that promotes the joint activity of pairs of neurons that belong to a certain assembly.

Assuming a constant external correlation matrix, the structure of the resulting self-connected assemblies of neurons can be quantified via the same graph theoretic measures as used previously (Fig. 8). We show that the tight assemblies previously observed for the modulation of the STDP and the EPSC functions (Fig. 5E and Fig. 7C) are formed for values of correlation strength one order of magnitude smaller than the synaptic upper bound. For the standard parameters of the minimal triplet model (Table 1; Fig. 7C, *η*_−_ = 1) for which the feedforward motif coefficients are the strongest (the motifs for which the *α*-path is longer than the *β*-path), we conclude that a significantly strong external correlation relative to internally generated network correlations is needed to promote common input.

**Figure 8.**
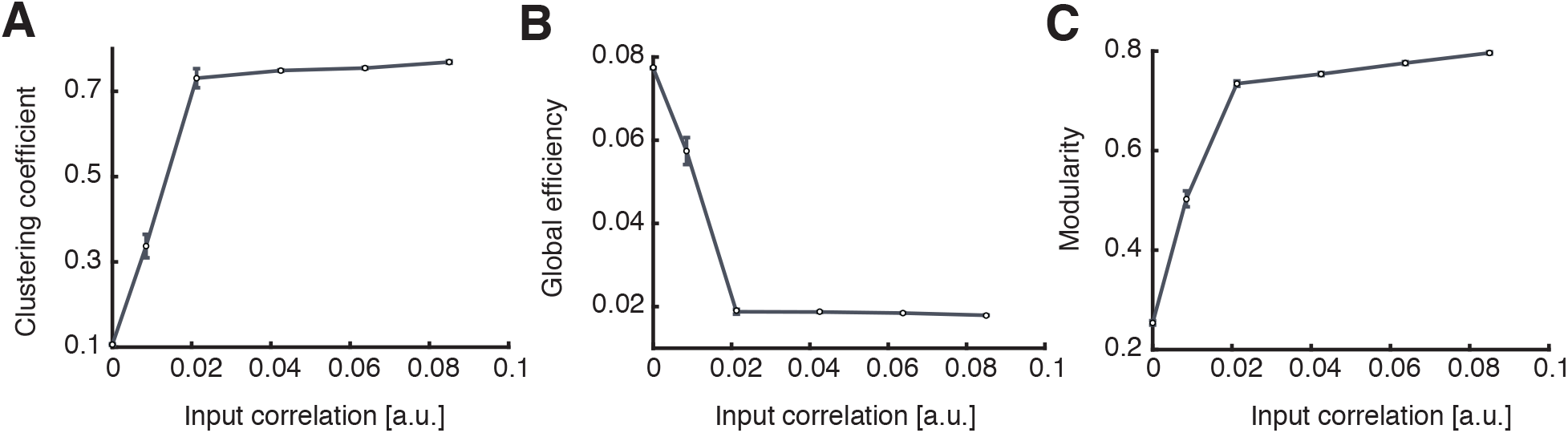
**A.** Mean clustering coefficient versus the pairwise correlation coefficient of the input pattern. The strength of the correlation was provided as ratios (0.01, 0.05, 0.125, 0.25, 0.375 and 0.5) of the possible maximum weight of each individual synaptic connection *w_max_*. **B.** Mean global efficiency versus the pairwise correlation coefficient of the input pattern. **C.** Modularity versus the pairwise correlation coefficient of the input pattern. The rapid increase of the clustering coefficient and the modularity, paired with a decrease of the global efficiency is a feature of robust assembly formation. Strong enough correlations in the external signal will generate tight assemblies.

## Discussion

We developed a self-consistent theoretical framework to study the impact of HOCs on the plasticity of recurrent networks by using the triplet STDP rule. Specifically, we derived the dependence of the drift in synaptic efficacy on network structure, taking into account contributions from structural motifs of all orders, and demonstrated the emergence of global network structures i.e. assemblies, from these local motifs. Based on recent work on the calculation of beyond second-order moments of mutually exciting Hawkes processes [37,99], we broke down the interactions including pairs and triplets of spikes to include paths of different length between neurons in the network along which information may propagate (Fig. 2 and 4). We characterized the unique motifs that arise from these higher-order interactions and analyzed their impact on the internal correlation structure and plasticity in the network through the motif coefficients (Fig. 3). While linearization of neuronal dynamics was required for this approach, it is a common technique used to approximate the dynamics of more realistic biophysical neurons [5,34]. Importantly, triplet interactions provide a way to include feedback loops in these recurrent networks as an alternative to using nonlinear neurons; feedback loops are not possible when linearizing networks with only pairwise interactions studied before [100].

We considered an asymmetric minimal triplet STDP rule, in which depression is induced by pairs of spikes and, conversely, potentiation is induced by triplets of spikes. This rule has been shown to describe plasticity experiments that the classical STDP rule, based on pairs of spikes, has failed to capture; for instance, plasticity experiments in which the pairing frequency during plasticity induction was varied [52,53]. As such, the triplet STDP rule is sensitive to HOCs. HOCs have not only been measured in the brain, but also shown to play an important role in visual coding and representing experimental data [55,56,101,102]. HOCs are ubiquitous in sensory stimuli, such as natural stimuli and speech signals [103,104]. These correlations have been previously utilized in learning rules, such as the BCM rule, to extract the independent components or features in natural images resulting in simple cell receptive fields as seen in V1 [103,105–107]. Because of its mapping to the BCM rule [74], we can interpret the triplet SDP rule as a method for performing similar computations.

Modulating either the STDP function (Fig. 5) or the EPSC function (Fig. 7) enabled the spontaneous formation of self-connected assemblies without the need for externally patterned inputs [47–49] or assuming a symmetric pair-based STDP rule [6]. We quantified the nature of the emergent assemblies using three graph-theoretic measures used to characterize spontaneous assemblies in the tectum of zebrafish larvae [25], offering insight into how modulating plasticity and synaptic transmission affects network structure in these experimental systems shaped by spontaneous activity. Interestingly, the final assemblies formed by modulating the EPSC function were more consistent across networks with different initial connectivity than the assemblies generated through the modification of the STDP function. This could be seen by the ‘tighter’ structures in the average connectivity matrices (Fig. 5E and Fig. 7C), and the higher values of modularity. The ultimate connectivity structure was determined by the relative strength of motifs which were regulated differently by each modulatory mechanism. In particular, modifying the EPSC function reinforced the influence of the common input motif (driven by the motif coefficient *M*_1,1_) over all others (Fig. 7B). In comparison, the modulation of the STDP rule maintained a strong feedforward drive (driven by the feedforward motif coefficients *M*_1,0_, *M*_2,1_, *M*_2,0_ and *M*_3,0_) and by making the corresponding reciprocal motif coefficients (*M*_0,1_, *M*_1,2_, *M*_0,2_ and *M*_0,3_) positive, reduced the competition between reciprocal connections. Together with maintaining a strong *M*_1,1_ coefficient, this generated robust assemblies (Fig. 5B). Applying external correlated input led to the emergence of self-organized assemblies that were similar to the assemblies from changing the EPSC function. Consequently, we propose that the mechanisms that promote the formation of assemblies can be diverse in different circuits depending on the nature of the plasticity rules, synaptic transmission (EPSC function) or the structure of external input that dominate in these circuits.

The action of different neuromodulators provide a way of modulating the STDP or synaptic transmission functions, and studying the effects of diverse neuromodulatory states on the plasticity of connections in many brain regions is possible with recent advances in experimental techniques [84–86]. For instance, timing properties of STDP have been shown to depend on the dynamics of the latent signaling pathways involving these factors. Understanding the consequences of changing the properties of the underlying plasticity mechanisms on network dynamics can further elucidate the process of learning and memory storage in recurrent networks found everywhere in the brain [84–86].

Our framework enabled us to derive specific connectivity structures that emerge in recurrent networks such as assemblies, which have been abundantly observed in experimental data. Connectivity matrices of large recurrent networks are generally difficult to assay experimentally, requiring multiple cells to be patched simultaneously [79], although recent developments in the field of connectomics offer potential for these matrices to be obtained in the future [108,109]. However, a good experimental determinate of assemblies may be derived from functional interactions among neurons, inferred from physiological experiments that simultaneously record the activity of a large number of neurons. While it is clear that neuronal activity exhibits structure in response to sensory input, assemblies are present even during spontaneous activity and have similar spatial organization [21,25,26]. This has suggested that these self-organized assemblies are biologically relevant for the processing of information in these networks and for the representation of sensory stimulus attributes [21]. In the rodent visual cortex, a given stimulus, of the form of a natural scene or an orientated grating, consistently activates a specific assembly [21]. Supporting evidence on the behavioral scale argues that the functional circuit connectivity may be intrinsically adapted to respond preferentially to stimuli of biological relevance for the survival of the animal, such as catching prey or avoiding predators [24,27].

Our analytical approach offers a precise description of how ongoing plasticity shapes connectivity in recurrent networks driven by spontaneous activity. Such spontaneous activity is especially common during early postnatal development, where it activates neural networks before the onset of sensory experience and the maturation of sensory organs. In the rodent visual system, for instance, eye opening only occurs during the second postnatal week of development [110]. Prior to this, spontaneous patterns of activity propagate throughout the entire visual system, including the retina, thalamus and cortex [111], which are known to instruct different aspects of circuit organization [112]. Interestingly, during very early postnatal development of somatosensory cortex in rodents (postnatal day 4), spontaneous activity exhibits a highly correlated state consisting of cell assemblies where multiple neurons show correlated activity [113]. By the second postnatal week this spontaneous activity transitions to a much more decorrelated state that lacks a clear spatial structure. A similar sparsification of spontaneous activity during development is also observed in the visual cortex, though lacking the spatial structure observed in the somatosensory cortex [114]. Since these two studies argue that over development functional connectivity becomes more desynchronized, this framework is more consistent with our analysis of the depression window of the STDP rule becoming smaller over development (Fig. 5). This broadening of the depression window in early development is consistent with a previously described burst-timing-dependent plasticity where the temporal integration of activity occurs over much longer timescales on the order of several hundred milliseconds than in adulthood [112,115,116].

Unlike individual neurons which acquire specific feature tuning before eye opening, for instance to orientation, functionally stronger connections between neurons with similar response properties occur only after eye opening [14]. This suggests that the emergence of functional specificity in recurrent connections among similarly tuned neurons in mouse primary visual cortex is driven by specific correlated input coming through the open eyes. Indeed, introducing higher-order spike interactions through a functionally similar plasticity rule to triplet STDP that we considered [89], enabled the formation of recurrent connection specificity observed experimentally [14]. More generally, assemblies have been shown to emerge in recurrent network models with balanced excitation and inhibition [47,48,117]. These assemblies exhibit attractor dynamics which have been argued to serve as the substrate of different computations, such as predictive coding through the spontaneous retrieval of evoked response patterns [47,48,117]. We investigated how structure emerges primarily from spontaneous dynamics in the case where HOCs drive synaptic plasticity through the triplet STDP rule. Considering these HOCs is key, as plasticity rule based on pairs of spike like the asymmetric pair-based STDP, cannot support the formation of self-connected assemblies [89], beyond promoting local reciprocal connections [91].

## Methods

### Network dynamics

We consider that the time dependent activity of a neuron *i* is given by a stochastic realization of an inhomogeneous Poisson process [67], with expectation value

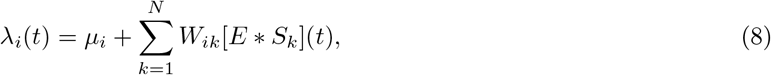

where *μ_i_* is the rate of spontaneous activity due to constant external input, **W** is the synaptic weight matrix, *S*(*t*) is the spike train and *E*(*t*) is the EPSC function, which we assume to be identical for all neurons. Then, the product **W***E*(*t*) is referred to as the interaction kernel. The operator ‘*’ corresponds to the convolution operation.

We assume that the overall effect of the inhibitory population on the synaptic efficacies among excitatory neurons is to balance the network activity. Thus, the sum of inhibitory synapses into each neuron is dynamically adjusted to match the sum of the excitatory synaptic efficacies, such that each element of the inhibitory connectivity matrix is equal to the average of the excitatory input as

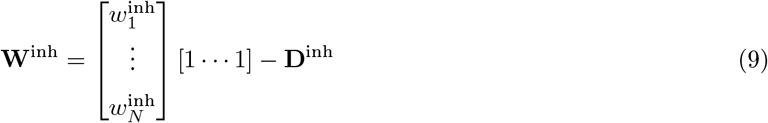

where

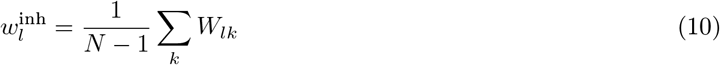

is the value of each row element and

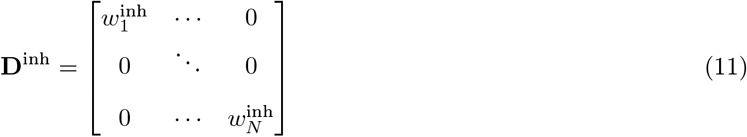

is a diagonal matrix to take into account there is no self-connectivity. The inhibitory connections are fast and updated in each integration step.

Plasticity of the connectivity matrix **W** is determined by pair-based and triplet STDP rules. We assume ‘all-to-all’ interactions between spikes, where each postsynaptic spike interacts with every previous pre- and postsynaptic spike and vice-versa [50,118–120].

We also implement heterosynaptic competition based on previous work [6, 78] as an additional mechanism for the plasticity dynamics to restrict the maximum number of strong connections a neuron can make, and thus keep the spectral radius of the connectivity matrix lower than one. The total synaptic input and output of each neuron is limited: the sum of the inbound (afferent) connections to each postsynaptic neuron *i* and the sum of outbound (efferent) excitatory synaptic efficacies from each presynaptic neuron *j* have an upper bound *W*_max_. The plasticity due to heterosynaptic competition can be written as

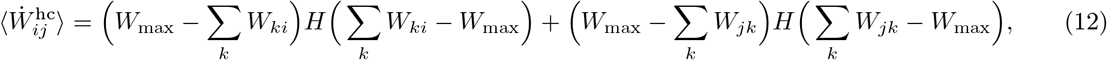

where *H* is the Heaviside function. Imposing an upper bound *w*_max_ for each synaptic efficacy restricts the possible number of connections a neuron can make to 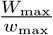. Therefore, the average amount of plasticity is the sum of the change due to STDP based on Eq. 2 and heterosynaptic competition based on Eq. 12.

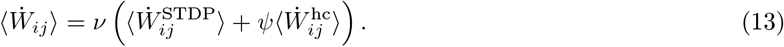

Here, the learning rate scale *ν* ensures that the synaptic efficacy increments in each integration step are small. The relative contribution of heterosynaptic competition to overall plasticity is determined by the heterosynaptic competition term *ψ*. The value for these parameters can be found in Table 1.

### Averaged synaptic efficacy dynamics for pair-based and triplet STDP rules

Assuming slow learning in comparison to neuronal dynamics and that pairs and triplets of spikes between the pre- and postsynaptic neurons are relevant to plasticity [52,74], the mean evolution of the synaptic efficacies due to STDP is given by

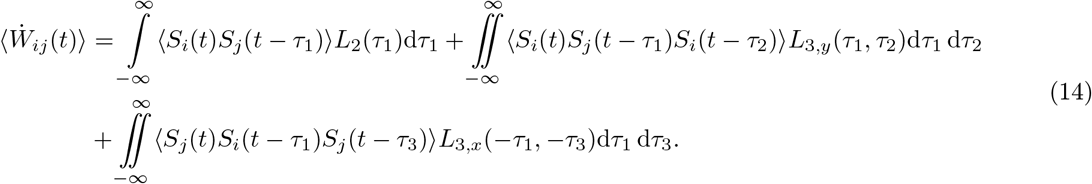

*L*_2_ corresponds to the pair-based STDP rule and *L*_3_ to the triplet STDP rule. The additional subscripts *x* and *y* denote that the triplets which contribute to plasticity are two pre- and one postsynaptic spikes and one pre- and two postsynaptic spikes, respectively. *τ*_1_ is the time difference between the spikes of the pre- and the postsynaptic neuron. *τ*_2_ is the time difference between two postsynaptic spikes and *τ*_3_ is the time difference between two presynaptic spikes (Fig. 1B). It should be highlighted that this derivation is independent of the specific shape of the STDP functions.

By definition, the mean rates of the pre- and postsynaptic neurons are *r_j_* and *r_i_* accordingly. We consider both to be stationary at equilibrium. The second-order correlation between the pre- and postsynaptic neurons with time delay *τ*_1_ is

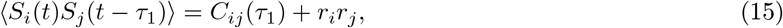

where *C_ij_* is an element of the covariance matrix (Fig. 1B). Note that [6,37] use a different convention for signs. The third-order correlation between the triplet of spikes ‘post-pre-post’ with *τ*_1_ and *τ*_2_ is

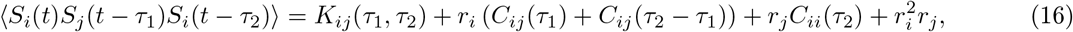

where *K_ij_* denotes a third-order cumulant [74]. Analogously, for the ‘pre-post-pre’ third-order correlation

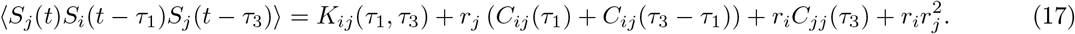

### Calculation of cumulants

The definition of cumulants in the Fourier space is imperative for the derivation of our results. Assuming stationarity, the expected firing rate (i.e. the first order cumulant) vector **r** = 〈λ(*t*)〉 no longer depends on time and can be written as

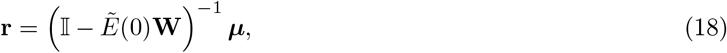

where 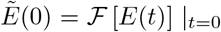 denotes the Fourier transform of the EPSC function evaluated at zero. For all the calculations, we define the Fourier transform as

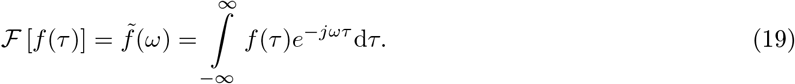

The second-order cumulant, the covariance, can be calculated in the time domain as [37, 99]

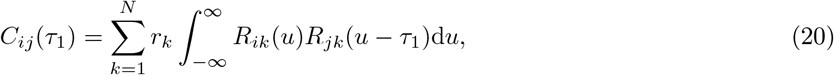

where 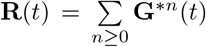 is defined as a ‘convolution power series’ [37,99] of the interaction kernel **G**(*t*) = **W***E*(*t*), with

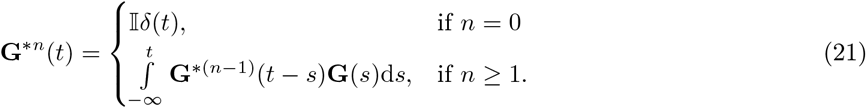

Formally, the computation of each element *R_mn_*(*t*) consists of calculating the probability of a spike from neuron *m* at time *t* given that neuron *n* fired at time 0. This representation provides a convenient formalism for representing causality of spiking events in our model. Then, considering the definition of ‘path lengths’ *α* and *β* from the source neuron *k* to the postsynaptic neuron *i* and the presynaptic neuron *j* (Fig. 2A), we can rewrite Eq. 20 as

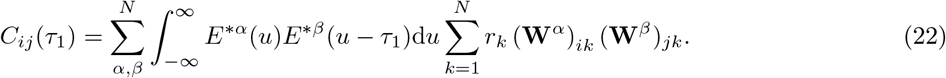

For the auto-covariance *C_ii_* for path lengths *α* and *γ* from the source neuron *k* to the postsynaptic neuron *i* (Fig. 2B) we analogously obtain

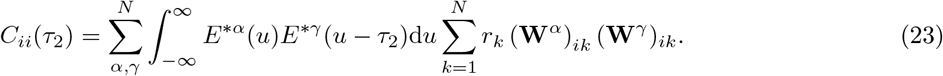

Since each *R* function consists of the convolution of the EPSC functions, then its Fourier transform is the product of the Fourier transforms of each of those functions, which simplifies calculations. Therefore, the cross-covariance *C_ij_* in the frequency domain (i.e. the Fourier transform of Eq. 20) is given by (detailed derivation in S1)

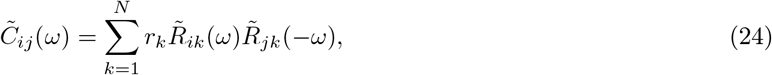

and, finally we obtain the expression

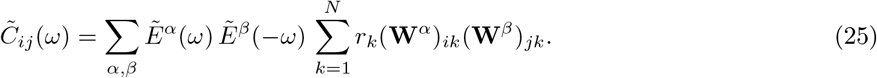

It should be noted that Eq. 25 was also derived in previous works using a different approach [6,67]. However, for the third-order cumulant *K_ij_* (Fig. 4) that same approach is not possible. Therefore, it is convenient to write *K_ij_* in the time domain in terms of the previously defined **R** [37,99] as

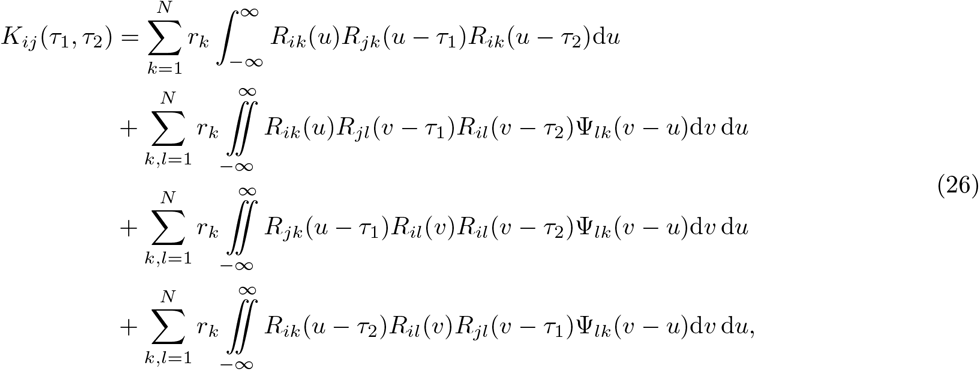

where additionally

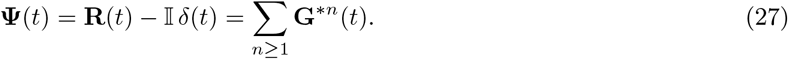

In Eq. 26, Ψ_*lk*_ (*v* – *u*) is the probability density of the event that a spike from neuron *k* at a time *v* – *u* = 0 causes a neuron *l* (different from neuron *k*) to emit a spike at a time *v* – *u* ≠ 0, after at least one synaptic connection. The function Ψ is necessary in Eq. 26 to take into account the branching structures in the calculation of *K_ij_* (Fig. 4B-D). In addition to *α*, *β* and *γ*, *ζ* is the path length from the source neuron *k* to the neuron *l* where the synaptic connection path branches out and is equal to or larger than one. Then, replacing both the *R* and Ψ functions by their corresponding definitions in terms of the connectivity matrix **W** and EPSC function *E*(*t*) yields

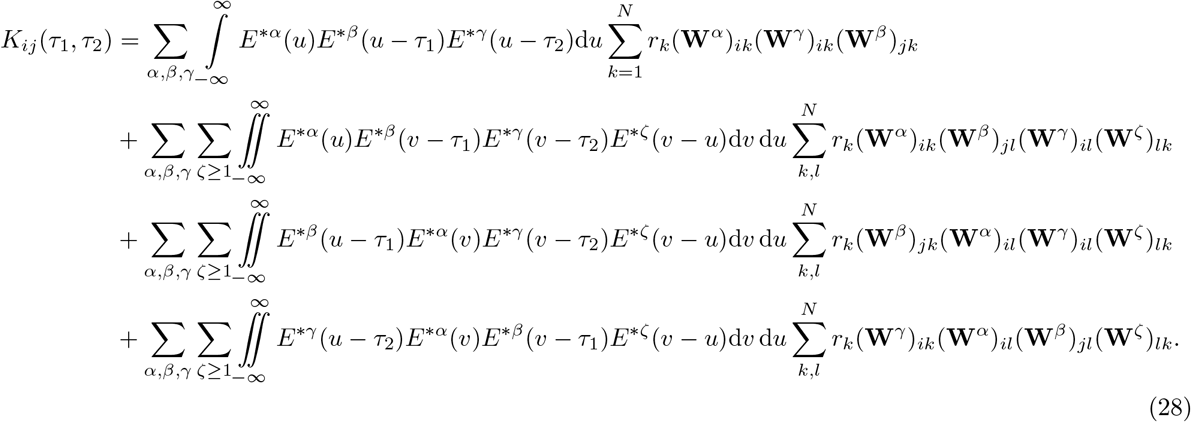

As with the covariance, we can calculate the Fourier transform of the third-order cumulant *K_ij_* from Eq. 26 as (detailed derivation in S1)

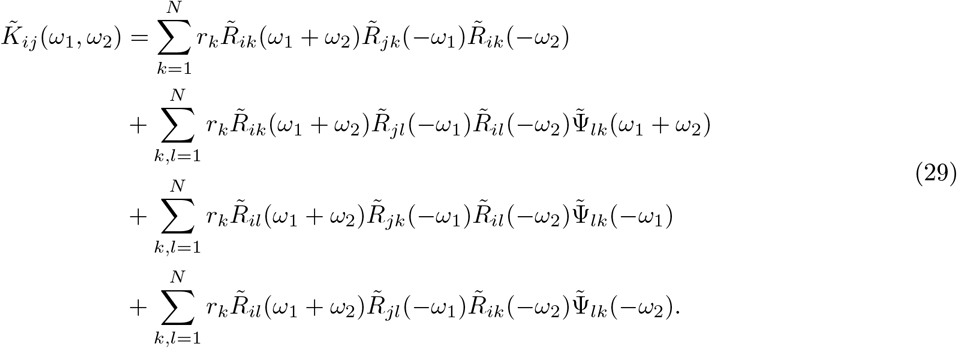

Finally, we obtain the third-order cumulant *K_ij_* in the Fourier domain in terms of the connectivity matrix **W**, the EPSC function *E*(*t*), and the path lengths *α*, *β*, *γ* and *ζ* as

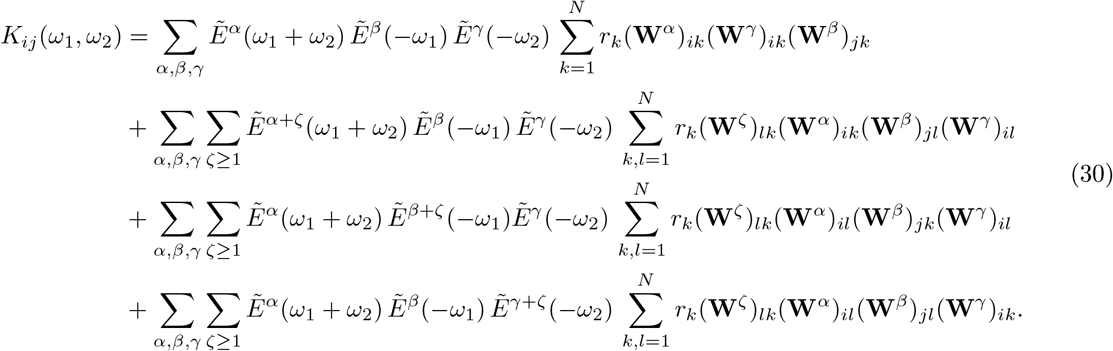

### Calculation of motif coefficients

To obtain the expression for the motif coefficients necessary for Eqs. 3, 4 and 5, we first need to insert Eqs. 25 and 30, i.e. the definitions of the second- and third-order cumulants in the frequency domain, in Eq. 2. Then, it can easily be seen that it is possible to separate the part that depends on the products of the connectivity matrix **W** from the rest. This way we define the motif coefficients as the integral of the products of the Fourier transforms of the STDP functions and the EPSC functions, considering the appropriate path lengths *α*, *β*, *γ* and *ζ* (Figs. 3B and D). In particular, for the motif coefficients in Eq. 3 we derive

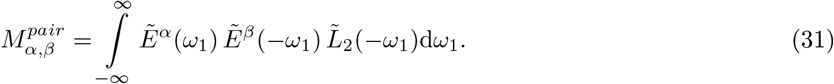

and

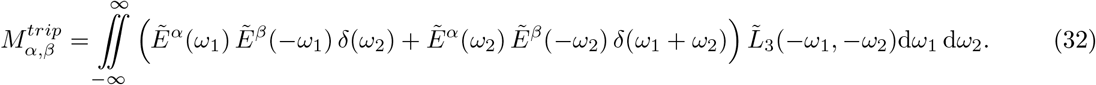

We note that this definition combines motif coefficients where *α* is the index corresponding to paths to the postsynaptic neuron, regardless of which of the two postsynaptic spike of the spike triplet it refers to (Fig. 1C). For the motif coefficient in Eq. 4 we derive

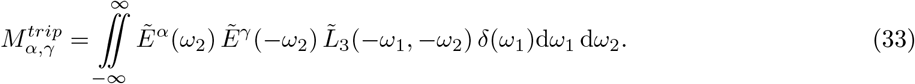

Lastly, for the ‘straight’ triplet motif (Fig. 4A) in Eq. 5 we get:

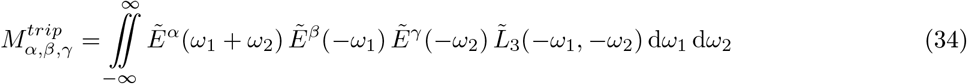

while for the ‘branching’ motifs (Fig. 4B-D) in Eq. 5:

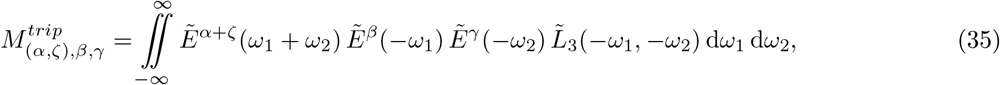

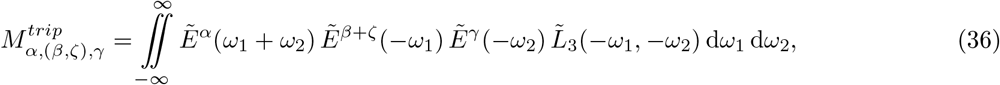

and

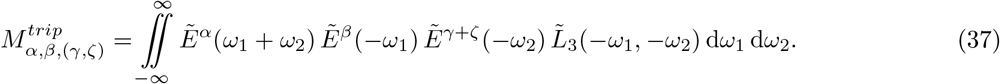

These expressions give us a concise representation of how the spiking activity interacts with network structure to impact plasticity.

### Synaptic dynamics

To calculate the values for the motif coefficients in Eqs. 31–37, we define the EPSC function *E*(*t*) as

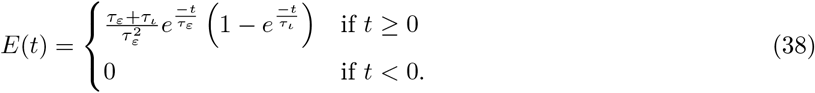

This function depends on two time constants *τ_ε_* and *τ_ι_* that define the onset and decay of the increase in the membrane potential with each spike. In particular, when *τ_ι_* → 0 the current is instantaneous and decays exponentially. The function is normalized to have an integral equal to 1, so that on average the number of postsynaptic spikes with the arrival of a presynaptic spike scales with the same order of magnitude as the synaptic efficacy. Its Fourier transform is

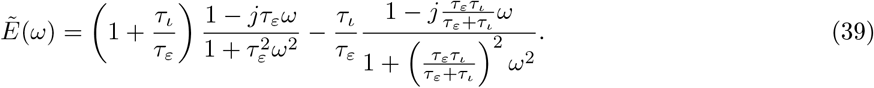

With respect to the choice of STDP function, we consider the minimal triplet STDP rule [52,74] that consists of the pair-based STDP function for depression and of a triplet STDP function for potentiation (Fig. 1C). Furthermore, we introduce a ‘modulation parameter’ *η*_−_ to model the reshaping of the depression window of the STDP function via modulatory effects.

The depression window of STDP the function can be written as

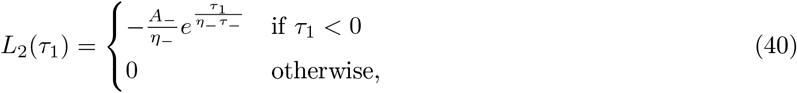

where *τ*_1_ = t_post_ – t_pre_ denotes the time difference between a post- and a presynaptic spike, *A*_−_ is the depression learning rate, *τ*_−_ is the depression time constant and *η*_−_ is the depression modulation parameter. The potentiation window of the STDP function depends on the timing of spike triplets (t_pre_, t_post_, 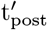)

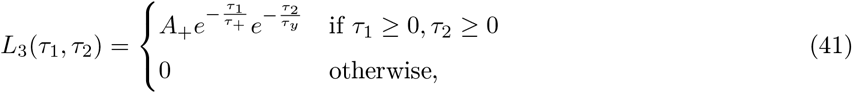

where again *τ*_1_ = t_post_ – t_pre_ denotes the time difference between a post- and a presynaptic spike and 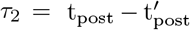 is the time difference between the two postsynaptic spikes; A_+_ is the potentiation learning rate, *τ*_+_ is the potentiation time constant and *τ_y_* is the second potentiation time constant.

While the ‘*A*’ parameters scale the amplitude of weight changes, the ‘*τ*’ coefficients determine how synchronous pre- and post-synaptic spikes must be to drive plasticity. The *η*_−_ parameter enables the modification of the shape of the STDP function. This additional parameter does not affect the total depression and potentiation in the rule and one can easily recover the ‘standard’ expressions for *η*_−_ → 1.

The Fourier transforms for these two functions are

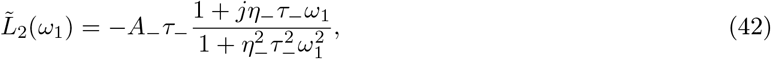

and

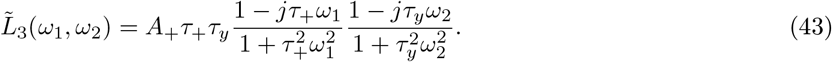

### Motif expansion up to third-order

After including Eqs. 3, 4 and 5 into Eq. 2, we can rewrite it in terms of the order of interactions in which they contribute to the averaged synaptic modification:

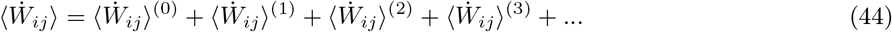

Assuming non-zero mean rates, no self-excitation (i.e. *W_ii_* = 0) and that terms of order higher than three can be disregarded in comparison to lower order ones, the terms of Eq. 44 are

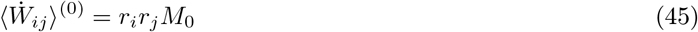

for the zeroth-order contributions,

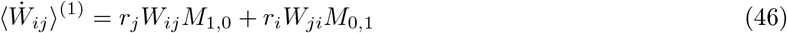

for the first-order contributions,

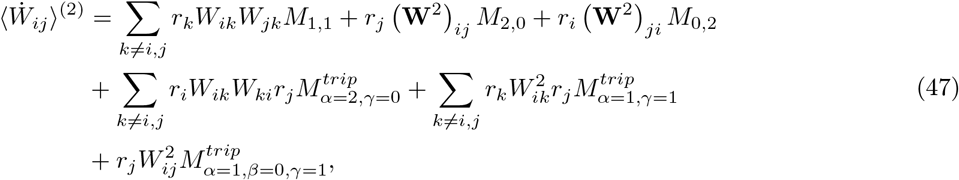

for the second-order contributions, and finally

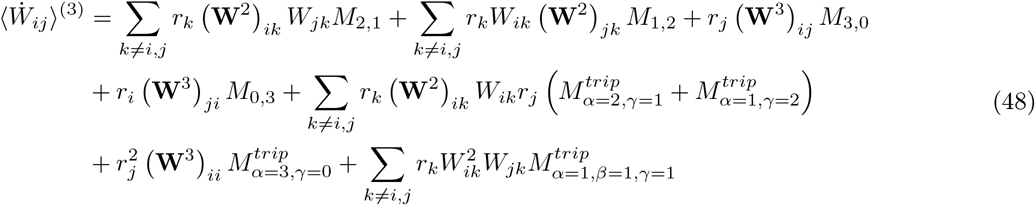

for the third-order contributions. Examples and illustrations of these motifs are given in Fig. 3. For conciseness, we grouped motif coefficients arising from the pair-based STDP rule and from the triplet STDP rule that shared values of *α* and *β* and relabeled them as

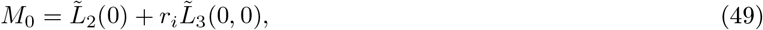

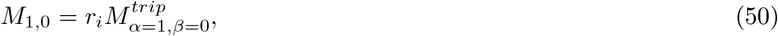

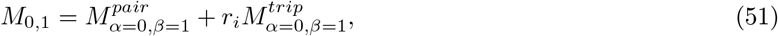

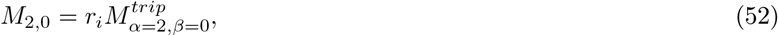

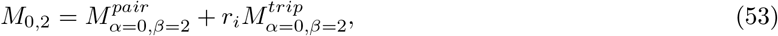

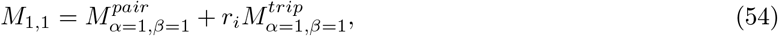

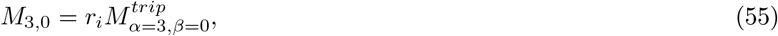

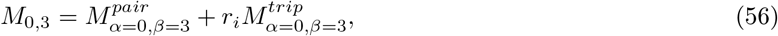

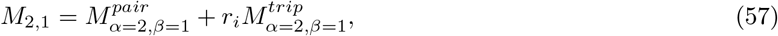

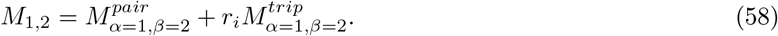

Using the defined functions for the EPSC (Eq. 39) and STDP functions (Eqs. 42 and 43) we calculated the motif coefficients in Eqs. 31–34, and consequently Eqs. 49–58. Note, we excluded the higher-than-third-order motif coefficients in Eqs. 35–37. These quantities represent the strength of contributions of each particular combination of paths from the source neuron to the pre- and postsynaptic neurons involved in the synaptic connection. Since we assume a learning rule balanced in potentiation and depression,

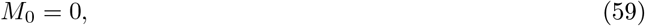

and thus

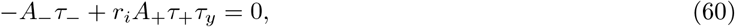

which is independent of the modulation parameter *η*_−_, this allows us to rewrite these motifs so that they are independent of the mean firing rate of the postsynaptic neuron. We analyze the evolution of these quantities in the main text, because they involve only *α* and *β* paths and remain constant throughout the numerical integration, in contrast to the motif coefficients in Eqs. 47 and 48 which involve both *α* and *γ* paths. The expressions for the motif coefficients defined by Eqs. 49–58 in terms of the EPSC and STDP functions’ parameters are given in S2.

### Averaged ordered connectivity matrices

The connectivity matrices resulting from integrating Eq. 2 numerically were ordered to reflect the graph structure of the network [6] (Figs. 5C). K-means classification was performed to group neurons that share similar connectivity, using a squared Euclidean distance. We then reordered the connectivity matrix based on the groups identified by the k-means clustering. Since the structures studied depend on initial conditions, despite the deterministic nature of our approach, we averaged the rearranged synaptic efficacy matrix over many trials with different (but random and weak) initial connectivity to obtain the most likely connectivity (Figs. 5E and 7C). Assemblies on the edges of the connectivity matrices have sharper edges due to an artifact created by the ordering algorithm, but this does not impact results.

### Network analysis

Graph theoretic measures for directed networks were calculated using algorithms of the Brain Connectivity Toolbox [121] from http://www.brain-connectivity-toolbox.net.

#### Clustering coefficient

For each connectivity matrix we computed the clustering coefficient [95]. For node *i*, this is

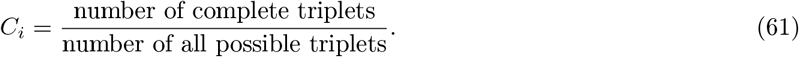

The number of complete triplets is obtained from the product of the corresponding edges of the node (from the adjancency matrix), and the total number of triplets depends on network size. Then, the average of the clustering coefficients of all the vertices *N* is given by [93]

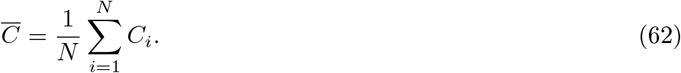

#### Global efficiency

The efficiency in the communication between nodes *i* and *j* can be defined to be inversely proportional to the shortest distance. The average efficiency of a network is calculated as [96]

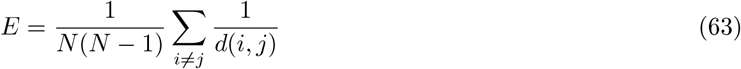

where *N* denotes the nodes in the network and *d*(*i*, *j*) is the length of the shortest path between a node *i* and a different node *j*. As an alternative to the average path length, the global efficiency of a network is defined as

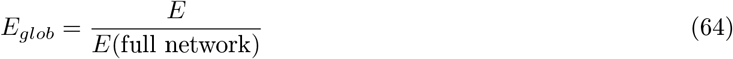

where the efficiency is scaled by an ideal graph where all the possible edges exist (i.e. full network). The difference between these measures is that the first measure quantifies the efficiency in a network where only one packet of information is being moved through it and the global measure quantifies the efficiency where all the vertices are exchanging packets of information with each other [96].

#### Modularity

The modularity *Q* of a connectivity matrix is a measure of the strength of its division into clusters or modules. Formally, modularity can be calculated as [94]

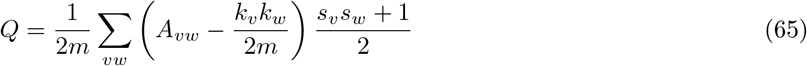

where **A** is the adjancency matrix of the graph, *k* is the node degree, *v* and *w* are the nodes’ indices and *s* is a variable that determines if the node belongs to a community or not. Modularity is the non-randomly distributed proportion of the edges that belong to the given cluster in a graph. It is positive if the number of edges within groups exceeds the number expected at random and depends on the chosen method for community detection.

The first algorithm we used for community detection, referred to as ‘spectral clustering’ algorithm, is based on the fact that modularity of a network is closely related to the structure of the eigenvalue spectrum of its weight matrix [94,122,123], high modularity means more strongly embedded communities. This is reflected in the spectra of the connectivity matrices as the separation of eigenvalues into a group with most eigenvalues and another of outliers, the number of which is often used to estimate the number of communities present in the network [94,122,123]. The second algorithm, called the Louvain method [98,121] is a greedy optimization method. First, smaller cliques are found by optimizing modularity locally on all nodes, then each small-sized community is grouped into one node and the first step is repeated. The complete modularity is then calculated by maximizing this value over all the divisions of the network into clusters [98,121]. We did not find any relevant differences between the Louvain method [98] and the spectral clustering algorithm [94,122,123], which were used to define community structure (see Fig. 6C).

## Supporting information

**S1. Fourier transform of the second- and third-order cumulants.**

**S2. Calculation of motif coefficients up to third-order.**

## Acknowledgments

LM and JG thank the Max Planck Society for funding. We thank members of the “Computation in Neural Circuits” group for useful discussions, Leonidas Richter for reading the manuscript and especially Christoph Miehl for critical feedback on the technical aspects of the manuscript. We thank Neta Ravid Tannenbaum and Yoram Burak for sharing the code used to obtain the connectivity matrices, and to Lilach Avitan and Jan Mölter for sharing further information regarding the methods used for community detection in their publications.

## References

1. Hebb DO. The Organization of Behavior. John Wiley & Sons; 1949.

2. Löwel S, Singer W. Selection of intrinsic horizontal connections in the visual cortex by correlated neuronal activity. Science. 1992;255(5041):209–212.

3. Shatz CJ. The Developing Brain. Scientific American. 1992;267(3):60–67.

4. Singer W. Neuronal representations, assemblies and temporal coherence. In: Progress in Brain Research. Elsevier; 1993. p. 461–474.

5. Ocker GK, Litwin-Kumar A, Doiron B. Self-organization of microcircuits in networks of spiking neurons with plastic synapses. PLoS Computational Biology. 2015;11(8):e1004458. doi:10.1371/journal.pcbi.1004458.

6. Ravid Tannenbaum N, Burak Y. Shaping neural circuits by high order synaptic interactions. PLoS Computational Biology. 2016;12(8):e1005056. doi:10.1371/journal.pcbi.1005056.

7. Carrillo-Reid L, Yang W, Bando Y, Peterka DS, Yuste R. Imprinting and recalling cortical ensembles. Science. 2016;353(6300):691–694. doi:10.1126/science.aaf7560.

8. Harris KD. Neural signatures of cell assembly organization. Nature Reviews Neuroscience. 2005;6(5):399–407. doi:10.1038/nrn1669.

9. Neves G, Cooke SF, Bliss TVP. Synaptic plasticity, memory and the hippocampus: a neural network approach to causality. Nature Reviews Neuroscience. 2008;9(1):65–75. doi:10.1038/nrn2303.

10. Lansner A. Associative memory models: from the cell-assembly theory to biophysically detailed cortex simulations. Trends in Neurosciences. 2009;32(3):178–186. doi:10.1016/j.tins.2008.12.002.

11. Buzsáki G. Neural syntax: Cell assemblies, synapsembles, and readers. Neuron. 2010;68(3):362–385. doi:10.1016/j.neuron.2010.09.023.

12. Perin R, Berger TK, Markram H. A synaptic organizing principle for cortical neuronal groups. Proceedings of the National Academy of Sciences. 2011;108(13):5419–5424. doi:10.1073/pnas.1016051108.

13. Ko H, Hofer SB, Pichler B, Buchanan KA, Sjöström PJ, Mrsic-Flogel TD. Functional specificity of local synaptic connections in neocortical networks. Nature. 2011;473(7345):87–91. doi:10.1038/nature09880.

14. Ko H, Cossell L, Baragli C, Antolik J, Clopath C, Hofer SB, et al. The emergence of functional microcircuits in visual cortex. Nature. 2013;496(7443):96–100. doi:10.1038/nature12015.

15. Holtmaat A, Caroni P. Functional and structural underpinnings of neuronal assembly formation in learning. Nature Neuroscience. 2016;19(12):1553–1562. doi:10.1038/nn.4418.

16. Yoshimura Y, Dantzker JLM, Callaway EM. Excitatory cortical neurons form fine-scale functional networks. Nature. 2005;433(7028):868–873. doi:10.1038/nature03252.

17. Kampa BM, Letzkus JJ, Stuart GJ. Cortical feed-forward networks for binding different streams of sensory information. Nature Neuroscience. 2006;9:1472–1473.

18. Miner D, Triesch J. Plasticity-driven self-organization under topological constraints accounts for non-random features of cortical synaptic wiring. PLoS Computational Biology. 2016;12(2):e1004759. doi:10.1371/journal.pcbi.1004759.

19. Cossell L, Iacaruso MF, Muir DR, Houlton R, Sader EN, Ko H, et al. Functional organization of excitatory synaptic strength in primary visual cortex. Nature. 2015;518:399–403. doi:10.1038/nature14182.

20. Lee WC, Bonin V, Reed M, Graham BJ, Hood G, Glattfelder K, et al. Anatomy and function of an excitatory network in the visual cortex. Nature. 2016;532:370–374. doi:10.1038/nature17192.

21. Miller JK, Ayzenshtat I, Carrillo-Reid L, Yuste R. Visual stimuli recruit intrinsically generated cortical ensembles. Proceedings of the National Academy of Sciences. 2014;111(38):E4053–E4061. doi:10.1073/pnas.1406077111.

22. Kruskal PB, Li L, MacLean JN. Circuit reactivation dynamically regulates synaptic plasticity in neocortex. Nature Communications. 2013;4(1). doi:10.1038/ncomms3574.

23. Gebhardt C, Baier H, Bene FD. Direction selectivity in the visual system of the zebrafish larva. Frontiers in Neural Circuits. 2013;7. doi:10.3389/fncir.2013.00111.

24. Romano SA, Pietri T, Pérez-Schuster V, Jouary A, Haudrechy M, Sumbre G. Spontaneous neuronal network dynamics reveal circuit’s functional adaptations for behavior. Neuron. 2015;85(5):1070–1085. doi:10.1016/j.neuron.2015.01.027.

25. Avitan L, Pujic Z, Mölter J, Poll MVD, Sun B, Teng H, et al. Spontaneous activity in the zebrafish tectum reorganizes over development and is influenced by visual experience. Current Biology. 2017;27(16):2407–2419.e4. doi:10.1016/j.cub.2017.06.056.

26. Pietri T, Romano SA, Pérez-Schuster V, Boulanger-Weill J, Candat V, Sumbre G. The emergence of the spatial structure of tectal spontaneous activity is independent of visual inputs. Cell Reports. 2017;19(5):939–948. doi:10.1016/j.celrep.2017.04.015.

27. Marachlian E, Avitan L, Goodhill GJ, Sumbre G. Principles of functional circuit connectivity: Insights from spontaneous activity in the zebrafish optic tectum. Frontiers in Neural Circuits. 2018;12. doi:10.3389/fncir.2018.00046.

28. Kriener B, Helias M, Aertsen A, Rotter S. Correlations in spiking neuronal networks with distance dependent connections. Journal of Computational Neuroscience. 2009;27(2):177–200. doi:10.1007/s10827-008-0135-1.

29. Pernice V, Staude B, Cardanobile S, Rotter S. How structure determines correlations in neuronal networks. PLoS Computational Biology. 2011;7(5):e1002059. doi:10.1371/journal.pcbi.1002059.

30. Zhao L, Beverlin B, Netoff T, Nykamp DQ. Synchronization from second order network connectivity statistics. Frontiers in Computational Neuroscience. 2011;5. doi:10.3389/fncom.2011.00028.

31. Roxin A. The role of degree distribution in shaping the dynamics in networks of sparsely connected spiking neurons. Frontiers in Computational Neuroscience. 2011;5. doi:10.3389/fncom.2011.00008.

32. Gaiteri C, Rubin JE. The Interaction of Intrinsic Dynamics and Network Topology in Determining Network Burst Synchrony. Frontiers in Computational Neuroscience. 2011;5. doi:10.3389/fncom.2011.00010.

33. Litwin-Kumar A, Doiron B. Slow dynamics and high variability in balanced cortical networks with clustered connections. Nature Neuroscience. 2012;15(11):1498–1505. doi:10.1038/nn.3220.

34. Trousdale J, Hu Y, Shea-Brown E, Josić K. Impact of network structure and cellular response on spike time correlations. PLoS Computational Biology. 2012;8(3):e1002408. doi:10.1371/journal.pcbi.1002408.

35. Pernice V, Deger M, Cardanobile S, Rotter S. The relevance of network micro-structure for neural dynamics. Frontiers in Computational Neuroscience. 2013;7. doi:10.3389/fncom.2013.00072.

36. Helias M, Tetzlaff T, Diesmann M. The correlation structure of local neuronal networks intrinsically results from recurrent dynamics. PLoS Computational Biology. 2014;10(1):e1003428. doi:10.1371/journal.pcbi.1003428.

37. Jovanović S, Rotter S. Interplay between graph topology and correlations of third order in spiking neuronal networks. PLoS Computational Biology. 2016;12(6):e1004963. doi:10.1371/journal.pcbi.1004963.

38. Hu Y, Trousdale J, Josić K, Shea-Brown E. Motif statistics and spike correlations in neuronal networks. Journal of Statistical Mechanics: Theory and Experiment. 2013;2013(03):P03012. doi:10.1088/1742-5468/2013/03/p03012.

39. Hu Y, Trousdale J, Josić K, Shea-Brown E. Local paths to global coherence: Cutting networks down to size. Physical Review E. 2014;89(3). doi:10.1103/physreve.89.032802.

40. Ocker GK, Hu Y, Buice MA, Doiron B, Josić K, Rosenbaum R, et al. From the statistics of connectivity to the statistics of spike times in neuronal networks. Current Opinion in Neurobiology. 2017;46:109–119. doi:10.1016/j.conb.2017.07.011.

41. Dechery JB, MacLean JN. Functional triplet motifs underlie accurate predictions of single-trial responses in populations of tuned and untuned V1 neurons. PLoS Computational Biology. 2018;14(5):e1006153. doi:10.1371/journal.pcbi.1006153.

42. Gilson M, Burkitt AN, Grayden DB, Thomas DA, van Hemmen JL. Emergence of network structure due to spike-timing-dependent plasticity in recurrent neuronal networks. I. Input selectivity–strengthening correlated input pathways. Biological Cybernetics. 2009;101(2):81–102. doi:10.1007/s00422-009-0319-4.

43. Gilson M, Burkitt AN, Grayden DB, Thomas DA, van Hemmen JL. Emergence of network structure due to spike-timing-dependent plasticity in recurrent neuronal networks. II. Input selectivity—symmetry breaking. Biological Cybernetics. 2009;101(2):103–114. doi:10.1007/s00422-009-0320-y.

44. Gilson M, Burkitt AN, Grayden DB, Thomas DA, van Hemmen JL. Emergence of network structure due to spike-timing-dependent plasticity in recurrent neuronal networks IV: structuring synaptic pathways among recurrent connections. Biological Cybernetics. 2009;101(5-6):427–444. doi:10.1007/s00422-009-0346-1.

45. Burkitt AN, Gilson M, van Hemmen JL. Spike-timing-dependent plasticity for neurons with recurrent connections. Biological Cybernetics. 2007;96:533–546. doi:10.1007/s00422-007-0148-2.

46. Triplett MA, Avitan L, Goodhill GJ. Emergence of spontaneous assembly activity in developing neural networks without afferent input. PLoS Computational Biology. 2018;14(9):e1006421. doi:10.1371/journal.pcbi.1006421.

47. Litwin-Kumar A, Doiron B. Formation and maintenance of neuronal assemblies through synaptic plasticity. Nature Communications. 2014;5(1). doi:10.1038/ncomms6319.

48. Zenke F, Agnes EJ, Gerstner W. Diverse synaptic plasticity mechanisms orchestrated to form and retrieve memories in spiking neural networks. Nature Communications. 2015;6(1). doi:10.1038/ncomms7922.

49. Ocker GK, Doiron B. Training and spontaneous reinforcement of neuronal assemblies by spike timing plasticity. Cerebral Cortex. 2018; doi:10.1093/cercor/bhy001.

50. Kempter R, Gerstner W, van Hemmen JL. Hebbian learning and spiking neurons. Physical Review E. 1999;59(4):4498–4514. doi:10.1103/physreve.59.4498.

51. Markram H. A history of spike-timing-dependent plasticity. Frontiers in Synaptic Neuroscience. 2011;3. doi:10.3389/fnsyn.2011.00004.

52. Pfister JP, Gerstner W. Triplets of spikes in a model of spike timing-dependent plasticity. Journal of Neuroscience. 2006;26(38):9673–9682. doi:10.1523/jneurosci.1425-06.2006.

53. Sjöström PJ, Turrigiano GG, Nelson SB. Rate, timing, and cooperativity jointly determine cortical synaptic plasticity. Neuron. 2001;32(6):1149–1164. doi:10.1016/s0896-6273(01)00542-6.

54. Wang HX, Gerkin RC, Nauen DW, Wang GQ. Coactivation and timing-dependent integration of synaptic potentiation and depression. Nat Neurosci. 2005;8:187–193.

55. Ohiorhenuan IE, Mechler F, Purpura KP, Schmid AM, Hu Q, Victor JD. Sparse coding and high-order correlations in fine-scale cortical networks. Nature. 2010;466(7306):617–621. doi:10.1038/nature09178.

56. Montani F, Ince RAA, Senatore R, Arabzadeh E, Diamond ME, Panzeri S. The impact of high-order interactions on the rate of synchronous discharge and information transmission in somatosensory cortex. Philosophical Transactions of the Royal Society A: Mathematical, Physical and Engineering Sciences. 2009;367(1901):3297–3310. doi:10.1098/rsta.2009.0082.

57. Ohiorhenuan IE, Victor JD. Information-geometric measure of 3-neuron firing patterns characterizes scale-dependence in cortical networks. Journal of Computational Neuroscience. 2010;30(1):125–141. doi:10.1007/s10827-010-0257-0.

58. Yu S, Yang H, Nakahara H, Santos GS, Nikolic D, Plenz D. Higher-order interactions characterized in cortical activity. Journal of Neuroscience. 2011;31(48):17514–17526. doi:10.1523/jneurosci.3127-11.2011.

59. Bohte SM, Spekreijse H, Roelfsema PR. The effects of pair-wise and higher order correlations on the firing rate of a post-synaptic neuron. Neural Comput. 2000;12(1):153–179.

60. Zylberberg J, Shea-Brown E. Input nonlinearities can shape beyond-pairwise correlations and improve information transmission by neural populations. Physical Review E. 2015;92(6). doi:10.1103/physreve.92.062707.

61. Barreiro AK, Gjorgjieva J, Rieke F, Shea-Brown E. When do microcircuits produce beyond-pairwise correlations? Frontiers in Computational Neuroscience. 2014;8. doi:10.3389/fncom.2014.00010.

62. Cayco-Gajic NA, Zylberberg J, Shea-Brown E. Triplet correlations among similarly tuned cells impact population coding. Frontiers in Computational Neuroscience. 2015;9. doi:10.3389/fncom.2015.00057.

63. Amari S, Nakahara H, Wu S, Sakai Y. Synchronous firing and higher-order interactions in neuron pool. Neural Computation. 2003;15(1):127–142. doi:10.1162/089976603321043720.

64. Macke JH, Opper M, Bethge M. Common input explains higher-order correlations and entropy in a simple model of neural population activity. Physical Review Letters. 2011;106(20). doi:10.1103/physrevlett.106.208102.

65. Montangie L, Montani F. Quantifying higher-order correlations in a neuronal pool. Physica A: Statistical Mechanics and its Applications. 2015;421:388–400. doi:10.1016/j.physa.2014.11.046.

66. Montangie L, Montani F. Common inputs in subthreshold membrane potential: The role of quiescent states in neuronal activity. Physical Review E. 2018;97(6). doi:10.1103/physreve.97.060302.

67. Hawkes AG. Spectra of some self-exciting and mutually exciting point processes. Biometrika. 1971;58(1):83–90. doi:10.1093/biomet/58.1.83.

68. Bi GQ, Poo MM. Synaptic modifications in cultured hippocampal neurons: dependence on spike timing, synaptic strength, and postsynaptic cell type. Journal of Neuroscience. 1998;18:10464–10472.

69. Brette R. Philosophy of the spike: Rate-based vs. spike-based theories of the brain. Frontiers in Systems Neuroscience. 2015;9. doi:10.3389/fnsys.2015.00151.

70. Gjorgjieva J, Drion G, Marder E. Computational implications of biophysical diversity and multiple timescales in neurons and synapses for circuit performance. Current Opinion in Neurobiology. 2016;37:44–52. doi:10.1016/j.conb.2015.12.008.

71. Hansel D, Mato G. Asynchronous states and the emergence of synchrony in large networks of interacting excitatory and inhibitory neurons. Neural Computation. 2003;15(1):1–56. doi:10.1162/089976603321043685.

72. Cessac B. On dynamics of integrate-and-fire neural networks with conductance based synapses. Frontiers in Computational Neuroscience. 2008;2:1–20. doi:10.3389/neuro.10.002.2008.

73. Renart A, de la Rocha J, Barthó P, Hollender L, Parga N, Reyes A, et al. The asynchronous state in cortical circuits. Science. 2010;327(5965):587–590. doi:10.1126/science.1179850.

74. Gjorgjieva J, Clopath C, Audet J, Pfister JP. A triplet spike-timing-dependent plasticity model generalizes the Bienenstock-Cooper-Munro rule to higher-order spatiotemporal correlations. Proceedings of the National Academy of Sciences. 2011;108(48):19383–19388. doi:10.1073/pnas.1105933108.

75. Vogels TP, Sprekeler H, Zenke F, Clopath C, Gerstner W. Inhibitory plasticity balances excitation and inhibition in sensory pathways and memory networks. Science. 2011;334(6062):1569–1573. doi:10.1126/science.1211095.

76. Luz Y, Shamir M. Balancing feed-forward excitation and inhibition via Hebbian inhibitory synaptic plasticity. PLoS Computational Biology. 2012;8(1):e1002334. doi:10.1371/journal.pcbi.1002334.

77. Rubin R, Abbott LF, Sompolinsky H. Balanced excitation and inhibition are required for high-capacity, noise-robust neuronal selectivity. Proceedings of the National Academy of Sciences. 2017;114(44):E9366–E9375. doi:10.1073/pnas.1705841114.

78. Fiete IR, Senn W, Wang CZH, Hahnloser RHR. Spike-time-dependent plasticity and heterosynaptic competition organize networks to produce long scale-free sequences of neural activity. Neuron. 2010;65(4):563–576. doi:10.1016/j.neuron.2010.02.003.

79. Song S, Sjöström PJ, Reigl M, Nelson S, Chklovskii DB. Highly nonrandom features of synaptic connectivity in local cortical circuits. PLoS Biology. 2005;3(10):e350. doi:10.1371/journal.pbio.0030350.

80. Yassin L, Benedetti BL, Jouhanneau JS, Wen JA, Poulet JFA, Barth AL. An embedded subnetwork of highly active neurons in the neocortex. Neuron. 2010;68(6):1043–1050. doi:10.1016/j.neuron.2010.11.029.

81. Field R, D’amour J, Tremblay R, Miehl C, Rudy B, Gjorgjieva J, et al. Heterosynaptic plasticity determines the set-point for cortical excitatory-inhibitory balance. bioRxiv. 2018; doi:10.1101/282012.

82. Froemke RC, Poo M, Dan Y. Spike-timing-dependent synaptic plasticity depends on dendritic location. Nature. 2005;434(7030):221–225. doi:10.1038/nature03366.

83. Cassenaer S, Laurent G. Conditional modulation of spike-timing-dependent plasticity for olfactory learning. Nature. 2012;482(7383):47–52. doi:10.1038/nature10776.

84. Frémaux N, Gerstner W. Neuromodulated spike-timing-dependent plasticity, and theory of three-factor learning rules. Frontiers in Neural Circuits. 2016;9. doi:10.3389/fncir.2015.00085.

85. Pedrosa V, Clopath C. The role of neuromodulators in cortical plasticity. A computational perspective. Frontiers in Synaptic Neuroscience. 2017;8. doi:10.3389/fnsyn.2016.00038.

86. Foncelle A, Mendes A, Jędrzejewska-Szmek J, Valtcheva S, Berry H, Blackwell KT, et al. Modulation of spike-timing dependent plasticity: Towards the inclusion of a third factor in computational models. Frontiers in Computational Neuroscience. 2018;12. doi:10.3389/fncom.2018.00049.

87. Bissière S, Humeau Y, Lüthi A. Dopamine gates LTP induction in lateral amygdala by suppressing feedforward inhibition. Nature Neuroscience. 2003;6(6):587–592. doi:10.1038/nn1058.

88. Seol GH, Ziburkus J, Huang S, Song L, Kim IT, Takamiya K, et al. Neuromodulators control the polarity of spike-timing-dependent synaptic plasticity. Neuron. 2007;55(6):919–929. doi:10.1016/j.neuron.2007.08.013.

89. Clopath C, Büsing L, Vasilaki E, Gerstner W. Connectivity reflects coding: a model of voltage-based STDP with homeostasis. Nature Neuroscience. 2010;13(3):344–352. doi:10.1038/nn.2479.

90. Gjorgjieva J, Evers JF, Eglen SJ. Homeostatic activity-dependent tuning of recurrent networks for robust propagation of activity. The Journal of Neuroscience. 2016;36(13):3722–3734. doi:10.1523/jneurosci.2511-15.2016.

91. Babadi B, Abbott LF. Pairwise analysis can account for network structures arising from Spike-Timing Dependent Plasticity. PLoS Computational Biology. 2013;9(2):e1002906. doi:10.1371/journal.pcbi.1002906.

92. Betzel R, Wood KC, Angeloni C, Geffen MN, Bassett DS. Stability of spontaneous, correlated activity in mouse auditory cortex. bioRxiv. 2018; doi:10.1101/491936.

93. Watts DJ, Strogatz SH. Collective dynamics of ‘small-world’ networks. Nature. 1998;393(6684):440–442. doi:10.1038/30918.

94. Newman MEJ. Modularity and community structure in networks. Proceedings of the National Academy of Sciences. 2006;103(23):8577–8582. doi:10.1073/pnas.0601602103.

95. Fagiolo G. Clustering in complex directed networks. Physical Review E. 2007;76(2). doi:10.1103/physreve.76.026107.

96. Latora V, Marchiori M. Efficient behavior of small-world networks. Physical Review Letters. 2001;87(19). doi:10.1103/physrevlett.87.198701.

97. Reichardt J, Bornholdt S. Statistical mechanics of community detection. Physical Review E. 2006;74(1). doi:10.1103/physreve.74.016110.

98. Blondel VD, Guillaume JL, Lambiotte R, Lefebvre E. Fast unfolding of communities in large networks. Journal of Statistical Mechanics: Theory and Experiment. 2008;2008(10):P10008. doi:10.1088/1742-5468/2008/10/p10008.

99. Jovanović S, Hertz J, Rotter S. Cumulants of Hawkes point processes. Physical Review E. 2015;91(4). doi:10.1103/physreve.91.042802.

100. Ocker GK, Josić K, Shea-Brown E, Buice MA. Linking structure and activity in nonlinear spiking networks. PLoS Computational Biology. 2017;13(6):e1005583. doi:10.1371/journal.pcbi.1005583.

101. Luczak A, Barthó P, Marguet SL, Buzsaki G, Harris KD. Sequential structure of neocortical spontaneous activity in vivo. Proceedings of the National Academy of Sciences. 2007;104(1):347–352. doi:10.1073/pnas.0605643104.

102. Pillow JW, Shlens J, Paninski L, Sher A, Litke AM, Chichilnisky EJ, et al. Spatio-temporal correlations and visual signalling in a complete neuronal population. Nature. 2008;454:995–999.

103. Olshausen BA, Field DJ. Emergence of simple-cell receptive field properties by learning a sparse code for natural images. Nature. 1996;381:607–609.

104. Simoncelli EP, Olhausen BA. Natural Image Statistics and Neural Representation. Annual Review in Neuroscience. 2001;24:1193–1216.

105. Bell AJ, Sejnowski TJ. An information-maximization approach to blind separation and blind deconvolution. Neural Comp. 1995;7:1129–1159.

106. Intrator N, Cooper LN. Objective function formulation of the BCM theory of visual cortical plasticity: Statistical connections, stability conditions. Neural Networks. 1992;5:3–17.

107. Blais BS, Intrator N, Shouval H, Cooper L. Receptive field formation in natural scene environments. Comparison of single-cell learning rules. Neural Comp. 1998;10:1797–1813.

108. Lichtman J, Helmstaedter M. Connectomics at the cutting edge: Challenges and opportunities in high-resolution brain mapping. Science. 2014;346(6209):651–651. doi:10.1126/science.346.6209.651-c.

109. Swanson LW, Lichtman JW. From Cajal to Connectome and Beyond. Annual Review of Neuroscience. 2016;39(1):197–216. doi:10.1146/annurev-neuro-071714-033954.

110. Blankenship AG, Feller MB. Mechanisms underlying spontaneous patterned activity in developing neural circuits. Nature Reviews Neuroscience. 2010;11:18–29.

111. Ackman JB, Burbridge TH, Crair MC. Retinal waves coordinate patterned activity throughout the developing visual system. Nature. 2012;490:219–225.

112. Richter LM, Gjorgjieva J. Understanding neural circuit development through theory and models. Current Opinion in Neurobiology. 2017;46:39–47. doi:10.1016/j.conb.2017.07.004.

113. Golshani P, Gonçalves JT, Khoshkhoo S, Mostany R, Smirnakis S, Portera-Cailliau C. Internally Mediated Developmental Desynchronization of Neocortical Network Activity. Journal of Neuroscience. 2009;29(35):10890–10899. doi:10.1523/jneurosci.2012-09.2009.

114. Rochefort NL, Garaschuk O, Milos RI, Narushima M, Marandi N, Pichler B, et al. Sparsification of neuronal activity in the visual cortex at eye-opening. Proceedings of the National Academy of Sciences. 2009;106(35):15049–15054. doi:10.1073/pnas.0907660106.

115. Butts DA, Kanold PO, Shatz CJ. A Burst-Based “Hebbian” Learning Rule at Retinogeniculate Synapses Links Retinal Waves to Activity-Dependent Refinement. PLoS Biology. 2007;5(3):e61. doi:10.1371/journal.pbio.0050061.

116. Winnubst J, Cheyne JE, Niculescu D, Lohmann C. Spontaneous Activity Drives Local Synaptic Plasticity In Vivo. Neuron. 2015;87:399–410. doi:10.1016/j.neuron.2015.06.029.

117. Miconi T, McKinstry JL, Edelman GM. Spontaneous emergence of fast attractor dynamics in a model of developing primary visual cortex. Nature Communications. 2016;7(1). doi:10.1038/ncomms13208.

118. Gerstner W, Kempter R, van Hemmen JL, Wagner H. A neuronal learning rule for sub-millisecond temporal coding. Nature. 1996;383(6595):76–78. doi:10.1038/383076a0.

119. Kistler WM, van Hemmen JL. Modeling synaptic plasticity in conjunction with the timing of pre-and postsynaptic action potentials. Neural Computation. 2000;12(2):385–405. doi:10.1162/089976600300015844.

120. Song S, Miller KD, Abbott LF. Competitive Hebbian learning through spike-timing-dependent synaptic plasticity. Nature Neuroscience. 2000;3(9):919–926. doi:10.1038/78829.

121. Rubinov M, Sporns O. Complex network measures of brain connectivity: Uses and interpretations. NeuroImage. 2010;52(3):1059–1069. doi:10.1016/j.neuroimage.2009.10.003.

122. Leicht EA, Newman MEJ. Community structure in directed networks. Physical Review Letters. 2008;100(11). doi:10.1103/physrevlett.100.118703.

123. Nadakuditi RR, Newman MEJ. Spectra of random graphs with arbitrary expected degrees. Phys Rev E. 2013;87:012803. doi:10.1103/PhysRevE.87.012803.

